# Structural Determinants of Phosphopeptide Binding to the N-Terminal Src Homology 2 Domain of the SHP2 Phosphatase

**DOI:** 10.1101/2020.03.27.012492

**Authors:** M. Anselmi, P. Calligari, J.S. Hub, M. Tartaglia, G. Bocchinfuso, L. Stella

## Abstract

SH2 domain-containing tyrosine phosphatase 2 (SHP2), encoded by *PTPN11*, plays a fundamental role in the modulation of several signaling pathways. Germline and somatic mutations in *PTPN11* are associated with different rare diseases and hematologic malignancies, and recent studies have individuated SHP2 as a central node in oncogenesis and cancer drug resistance. SHP2 structure includes two Src homology 2 domains (N-SH2 and C-SH2) followed by a catalytic protein tyrosine phosphatase (PTP) domain. Under basal conditions, the N-SH2 domain blocks the active site, inhibiting phosphatase activity. Association of the N-SH2 domain with binding partners containing short amino acid motifs comprising a phosphotyrosine residue (pY) leads to N-SH2/PTP dissociation and SHP2 activation. Considering the relevance of SHP2 in signaling and disease and the central role of the N-SH2 domain in its allosteric regulation mechanism, we performed microsecond-long molecular dynamics simulations of the N-SH2 domain complexed to 12 different peptides, to define the structural and dynamical features determining the binding affinity and specificity of the domain. Phosphopeptide residues at position −2 to +5, with respect to pY, have significant interactions with the SH2 domain. In addition to the strong interaction of the pY residue with its conserved binding pocket, the complex is stabilized hydrophobically by insertion of residues +1, +3 and +5 in an apolar groove of the domain, and interaction of residue −2 with both the pY and a protein surface residue. Additional interactions are provided by hydrogen bonds formed by the backbone of residues −1, +1, +2 and +4. Finally, negatively charged residues at position +2 and +4 are involved in electrostatic interactions with two lysines (Lys89 and Lys91) specific of the SHP2 N-SH2 domain. Interestingly, the MD simulations illustrated a previously undescribed conformational flexibility of the domain, involving the core β-sheet and the loop that closes the pY binding pocket.

## INTRODUCTION

### SH2 domains

The idea of protein modularity, with independently folding domains of conserved sequences, began with the discovery of Src homology 2 (SH2) domains [Mayer 2017]. Their name comes from the identification of sequences of ∼100 amino acids conserved in numerous cytosolic tyrosine kinases, including Src, and the appendix “2” indicates that this module is the second in the Src sequence [Yaffe 2002]. Today we know that the human genome codes for 121 SH2 domains, contained in 111 distinct proteins [Liu 2006, 2012]. The primary biochemical function of SH2 domains is to selectively recognize polypeptides containing a phosphotyrosine (pY), along with specific contiguous residues.

Tyrosine phosphorylation contributes to only ∼0.5% of the total phosphoproteome, yet it plays critical roles in eukaryotic cell regulation [Gopalasingam 2015]. Substrate specificities of kinases and phosphatases are broad, and their effects in signaling are controlled also by their location. The presence in their structures of domains devoted to protein-protein interactions leads to proper positioning of these enzymes close to their substrates [Pawson 1997]. In pY signaling, kinases “write” the phosphorylation signal, which can be “erased” by phosphatases. SH2 domains “read” this information, using it to localize signaling proteins correctly [Kaneko 2012]. As a general scheme, binding of an extracellular ligand to a receptor tyrosine kinase induces activation of the receptor, which phosphorylates itself and other nearby proteins. These phosphorylated tyrosine residues then function as docking sites for the SH2 domains of other proteins, which are thus recruited to the cell membrane or activated, causing propagation of the signal [Bradshaw 2002]. In addition, SH2 domains enhance tyrosine phosphorylation *in vivo* by protecting binding sites in their target proteins from dephosphorylation [Jadwin 2018].

### Structure of the SH2 domains

300 three-dimensional structures of approximately 70 different SH2 domains have been determined. They reveal a highly conserved topology [Gopalasingam 2015; Liu 2017]. These domains contain approximately 100 amino acids, with a central β strand, flanked by two α helices. These secondary structure elements are labeled according to their position along the sequence: βA αA βB βC βD βE βF αB βG (Figure 1A). Each residue is then numbered consecutively within the secondary structures [Eck 1993]. The central β-sheet divides the domain into two functionally distinct sides. The N-terminal side, flanked by helix αA, comprises the conserved pY binding pocket (formed by the BC loop); the C-terminal side, flanked by helix αB and the EF and BG loops, provides a more variable binding surface (specificity determining region) that typically engages residues C-terminal to the pY (Figure 1B) [Kuriyan 1997; Bradshaw 2002; Liu 2006,]. The structural arrangement of the domain complexes described above corresponds to the two requirements of SH2 domains: they must bind other proteins only when they are phosphorylated and they must associate specifically only to certain sequences.

**Figure 1.**
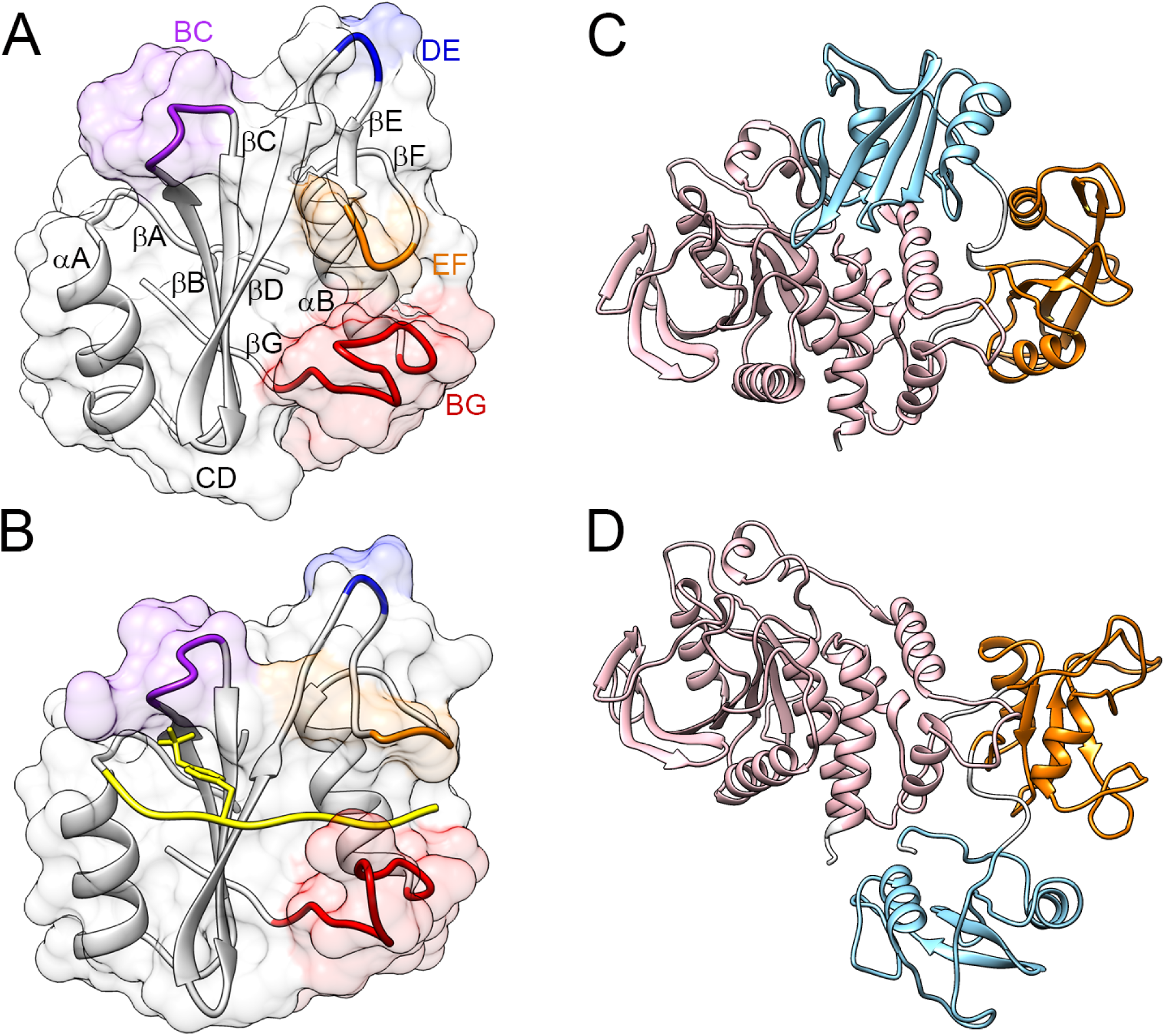
Structure of SHP2: N-SH2 domain and whole protein. The structure of the N-SH2 domain of SHP2 has the βαβββββαβ topology typical of SH2 domains (A). Loop BC (purple) is part of the pY binding pocket, loop DE (blue) inserts in the PTP active site in the autoinhibited SHP2 conformation, loops EF (orange) and BG (red) control access to the groove where the phosphopeptide binds. The crystallographic structures of the N-SH2 domain in the autoinhibited conformation of SHP2 (A) and when bound to a phosphopeptide (B) differ mainly for a rearrangement of the EF loop, which in the autoinhibited state blocks the peptide binding site of the N-SH2 domain. SHP2 comprises three domains: N-SH2 (light blue), C-SH2 (orange) and PTP (pink). In the absence of external stimuli, the N-SH2 domain blocks the catalytic site of the PTP domain (C). Binding of the SH2 domain to phosphorylated sequences, or pathogenic mutations, favor a conformational transition leading to a rearrangement of the domains and to activation (D). The structures in Panels C and D are reported with the same orientation of the PTP domain. PDB codes: 2SHP (A, C), 1AYA (B), 6CFR (D).

In most structures of SH2-ligand complexes, the phophopeptide binds across the surface of the domain, orthogonal to the central beta sheet, in an extended conformation (Figure 1B) [Kuriyan 1997], consistently with the observation that SH2 domains are able to associate to their cognate proteins even when these are denatured [Mayer 1998].

### The SH2 domain-containing protein tyrosine phosphatase 2

SH2 domains not only serve to connect the various components of signaling pathways by protein-protein interactions, but often also have a role in modulating enzymatic function. The SH2 domain-containing protein tyrosine phosphatases (PTPs) SHP1 and SHP2 contain two SH2 domains that are N-terminal to the catalytic domain, termed N-SH2 and C-SH2 (Figure 1C). In the absence of external stimuli, the N-SH2 domain interacts with the PTP active site, blocking it [Hof 1998]. Association of the SH2 domains to pY motifs favors N-SH2/PTP dissociation and thereby activation of the phosphatase (Figure 1D) [Gopalasingam 2015]. The loss of the N-SH2/PTP interactions is triggered by a conformational transition of N-SH2 that leads to a loss of complementarity between the N-SH2 and PTP surfaces.

The SHP2 protein was the first oncogenic PTP discovered. Mutations of *PTPN11* (the gene coding for SHP2) cause more than 30% of cases of juvenile myelomonocytic leukemia (JMML) [Tartaglia 2003; 2004a] and are variably found in other childhood malignancies [Tartaglia 2003, 2004b; Grossmann 2010]. In addition, SHP2 is required for survival of receptor tyrosine kinases (RTK)-driven cancer cells [Chen 2016], plays an important role in resistance to targeted cancer drugs [Ahmed 2019], is a mediator of immune checkpoint pathways [Okazaki 2013] and is involved in the induction of gastric carcinoma by *H. pylori* [Hayashi 2017]. *PTPN11* mutations also cause Noonan syndrome and Noonan syndrome with multiple lentigines, two disorders belonging to a family of rare diseases collectively known as RASopathies [Tartaglia 2001; Tartaglia 2004a; Tartaglia 2010]. For all these reasons, SHP2 is an important molecular target for therapies against cancer and rare diseases [Butterworth 2014; Ran 2016, Frankson 2017]. At the molecular level, pathogenic mutations of *PTPN11* often cause an increase in the binding affinity of the SH2 domains of SHP2, leading to hyper-activated signaling of the Ras/MAPK pathway [Tartaglia 2006; Bocchinfuso 2007; Martinelli 2008; Martinelli 2012].

Due to their role in many signaling pathways, SH2 domains have received much attention as potential targets of pharmaceuticals [Bradshaw 2002]. The fact that short pY-containing peptides (usually five to six amino acids) are sufficient to compete with larger protein ligands for SH2 domain binding has prompted researchers both in academia and industry to develop inhibitors of clinically relevant SH2 domains [Machida 2005]. However, no molecules targeting the SH2 domains of SHP2 for therapeutic purposes have been reported. Considering its role in the allosteric regulation of SHP2, the N-SH2 domain is particularly interesting under this respect.

### Phosphopeptide sequence selectivity of the N-SH2 domain of SHP2

Several proteins interacting with SHP2 through its SH2 domains have been identified. Lists of more than 50 known or putative interacting proteins have been compiled in the past [Songyang 1993; Sweeney 2005; Ihmof 2006] and several additional partners have been reported since then [Yang 2010; Kumamaru 2011; Tsutsumi 2013; Gandji 2015; Breitkopf 2016; Müller 2013; Vemulapalli 2019]. A database of the known interactions is available at phospho.elm.eu.org. However, in many of these cases, the sites of interaction, the pY residues that bind specifically to the SHP2 N-SH2 domain and the binding affinities have not been determined. Table 1 summarizes phosphorylated sequences for which a high binding affinity to the N-SH2 domain of SHP2 has been reported. Although exceptions do exist, a general consensus pattern can be clearly detected, with hydrophobic residues (A, L, I, V, M, F, P) at positions −2, +1, +3, +5, and acidic amino acids (D or E) at positions 2 and 4.

**Table 1.**
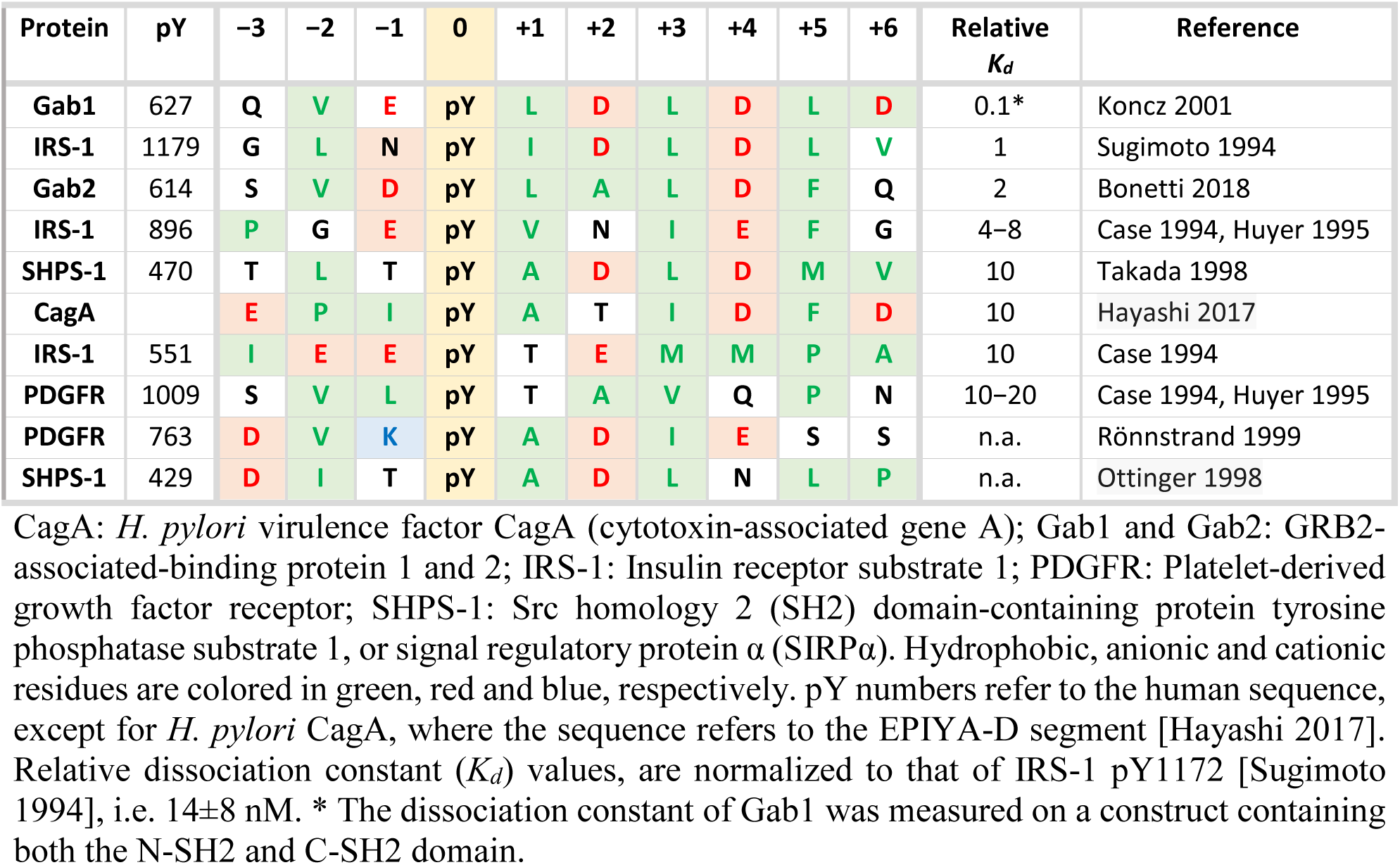
Natural Sequences with a High Affinity for the N-SH2 Domain of SHP2

The sequence selectivity of the N-SH2 domain of SHP2 has been analyzed also by utilizing phosphopeptide libraries. Oriented peptide library studies have examined positions from −1 to +6 with respect to pY. More recently, high throughput studies with surface immobilized peptide arrays [Huang 2008; Martinelli 2008, 2012; Tinti 2013] analyzed positions from −6 to +6, but distinct preferences were observed only in the −3 to +5 sequence stretch. The results of these investigations are summarized in Table 2. Collectively, a distinct preference for hydrophobic residues at positions −2, +1, +3 and +5 emerges (consistently with the natural sequences listed in Table 1), while other positions appear to be more variable. In particular, only peptide arrays indicated a possible preference for anionic residues in position +2 and +4.

**Table 2.** Motifs Determined from Peptide Library Studies

| -3 | -2 | -1 | 0 | +1 | +2 | +3 | +4 | +5 | +6 | Reference |
| --- | --- | --- | --- | --- | --- | --- | --- | --- | --- | --- |
|  |  |  | pY | IV |  | VILP |  |  |  | Songyang 1993 |
|  |  |  | pY | AVIT |  | LIV |  | WF |  | De Souza 2002 |
|  | ILV |  | pY | TVA |  | IVL |  |  |  | Sweeney 2005, I |
|  | W | XT | pY |  | IL |  |  |  |  | Sweeney 2005, II |
|  | IV |  | pY | LXT | Y | APTS |  |  |  | Sweeney 2005, III |
|  | IVL |  | pY | XF |  | P |  |  |  | Sweeney 2005, IV |
|  |  |  | pY |  |  |  | WYMH | FHWM | GPR | Imhof 2006 |
|  | LVH | LEIH | pY | VLIE | ED | ELMF | ELM |  |  | Huang 2008 |
|  | LV |  | pY | ILVA | NAD | LPV |  |  |  | Martinelli 2008 |
|  | LV | HT | pY | LIA | NA | LP | D | F |  | Martinelli 2012 |
| L | LVI | H | pY | LI | N | LP |  | F |  | Tinti 2013 |

X=norleucine. Sequence position investigated in each study are bordered in purple. Hydrophobic, anionic and cationic residues are colored in green, red and blue, respectively. Positions where one of these amino acid classes only is preferred are highlight.

Distinct selectivity features emerge from these data. Defining the determinants of N-SH2 selectivity is essential to allow the design of new peptides, peptidomimetics and small molecules targeted to this domain. To this end, we analyzed collectively the available X-ray structures and performed several molecular dynamics (MD) simulations of N-SH2/phosphopeptide complexes.

### Structures N-SH2/phosphopeptide complexes and MD simulations

Seven experimental structures of N-SH2/phosphopeptide complexes, obtained by X-ray crystallography, are available (PDB codes, 3TKZ, 3TL0, 4QSY, 1AYA, 1AYB, 1AYC, 5DF6, 5X7B, 5X94). In this work, 3TKZ and 1AYC were excluded from further analysis as in 3TKZ, a non-canonical 1:2 protein-peptide complex is formed (Zhang, 2011), while in 1AYC the N-SH2 domain is complexed with a nonspecific peptide (Lee, 1994). The phosphopeptides present in the remaining structures are listed in Table 3, which include the natural sequences of IRS-1 pY896 (1AYB), PDGFR pY1009 (1AYA), CagA (5X94 and 5X7B), Gab1 pY627 (4QSY).

**Table 3.**
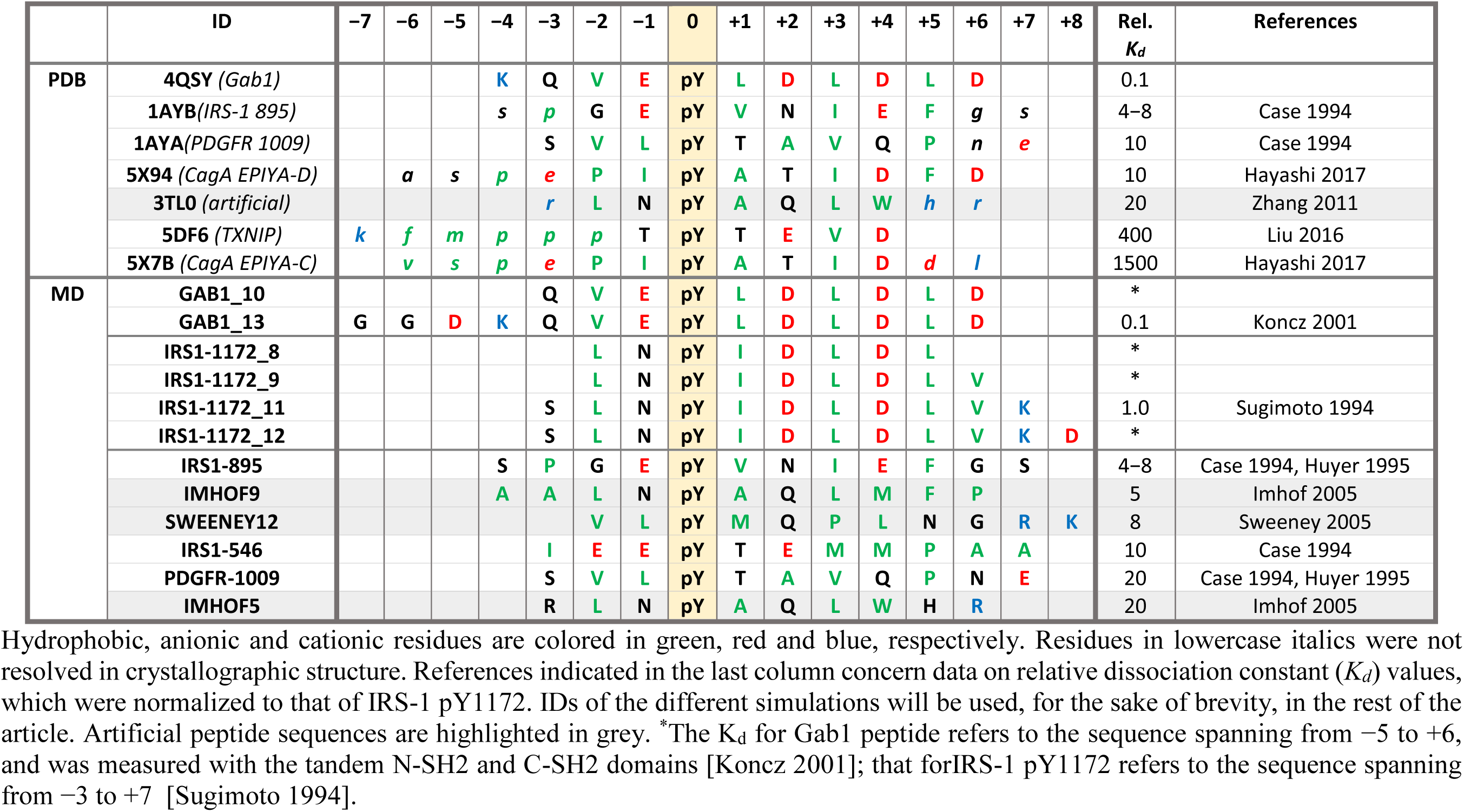
N-SH2/Peptide Complexes (Experimental and Simulated).

While these structures provide insights into the determinants of N-SH2 selectivity, characterization of the dynamics of domain/peptide complexes is essential to evaluate (i) the stability of the interactions observed in the crystallographic data and (ii) possible conformational transitions of the peptide or of the domain. In addition, no structures are available for the IRS-1 pY1179 peptide (which has one of the highest affinities among known sequences) or for high-affinity artificial peptides that were isolated in library screening studies. To address these issues, we performed twelve (microsecond-long) MD simulations of complexes of the N-SH2 domain with the Gab1, IRS-1 pY1172, pY895, pY546 (rat sequence numbering, corresponding to human pY1179, pY896 and pY551), PDGFR pY1009 and three artificial peptides isolated in [Sweeney 2005] and [Imhof 2006]. Moreover, for Gab1 and IRS-1 pY1172 we simulated several analogues of different lengths (Table 3).

## METHODS

Initial atomic coordinates were taken from crystallographic structures (Table S1). For some peptides (e.g. IRS1-895) the sequence in the simulation exactly matched the original sequence, and the crystal structure has been used as is. For other peptides (e.g. IRS1-1172 group), the crystal structure with the highest peptide sequence similarity was chosen, and the structure was edited by means of Molecular Operative Environment (MOE) (Chemical Computing Group, Inc.), followed by a conformational analysis and a local energy minimization with side chain repacking, yielding a reasonable binding pose for all peptides (Table S1). The termini of the peptides were capped by acetyl and amide groups. In all cases, the N-SH2 domain comprised residues 3 to 103. Each protein molecule was put at the center of a dodecahedron box, large enough to contain the domain and at least 0.9 nm of solvent on all sides. The protein was solvated with explicit TIP3P water molecules [Jorgensen 1983]. All MD simulations were performed with the GROMACS software package [Van Der Spoel 2005] using AMBER99SB force field [Hornak 2006] augmented with parm99 data set for phosphotyrosine [Homeyer 2006]. Long range electrostatic interactions were calculated with the particle-mesh Ewald (PME) approach [Darden 1993]. A cut-off of 1.2 nm was applied to the direct-space Coulomb and Lennard-Jones interactions. Bond lengths and angles of water molecules were constrained with the SETTLE algorithm [Miyamoto 1992], and all other bonds were constrained with LINCS [Hess 1997]. The pressure was set to 1 bar using the weak-coupling barostat [Berendsen 1984]. Temperature was fixed at 300 K using velocity rescaling with a stochastic term [Bussi 2007]. For all systems, the solvent was relaxed by energy minimization followed by 100 ps of MD at 300 K, while restraining protein atomic positions with a harmonic potential. The systems were then minimized without restraints and their temperature brought to 300 K in 10 ns in a stepwise manner. Finally, productive runs of 1 μs were performed.

Analysis of structural properties was performed using the GROMACS analysis tools. For crystallographic structures, hydrogen bonds were detected following the usual geometric criteria [Mills 1996].

## RESULTS AND DISCUSSION

### The −2 to +5 phosphopeptide region interacts tightly with the domain

During all simulations, peptides remained in the binding cleft for the whole length of the trajectory. Figure 2 reports the root mean square fluctuations (RMSF) of the position of phosphopeptide atoms and the order parameters of the side-chain Cα-Cβ bonds, calculated during the 12 MD simulations. In all cases, RMSF values were less than 1 Å for residues in the 0 to +4 interval, indicating a very low mobility for these peptide stretches. Consistently, order parameters were generally higher than 0.75 in this peptide region, although some exceptions were present at positions +1 and +4. In many cases, also residues −2, −1 and +5 were rather stable, although a larger variability was observed compared to the central stretch. The structures of Figure 2 show that the peptide termini (out of the −2 to +5 region) can detach from the protein. Overall, these findings explain why a distinct selectivity was observed in the peptide library studies only for amino acids falling in the interval from −2 to +5 (Tables 1 and 2). This conclusion is supported by the fact that residues preceding −2 or following +5 are often unresolved in X-ray structures (Table 3).

**Figure 2.**
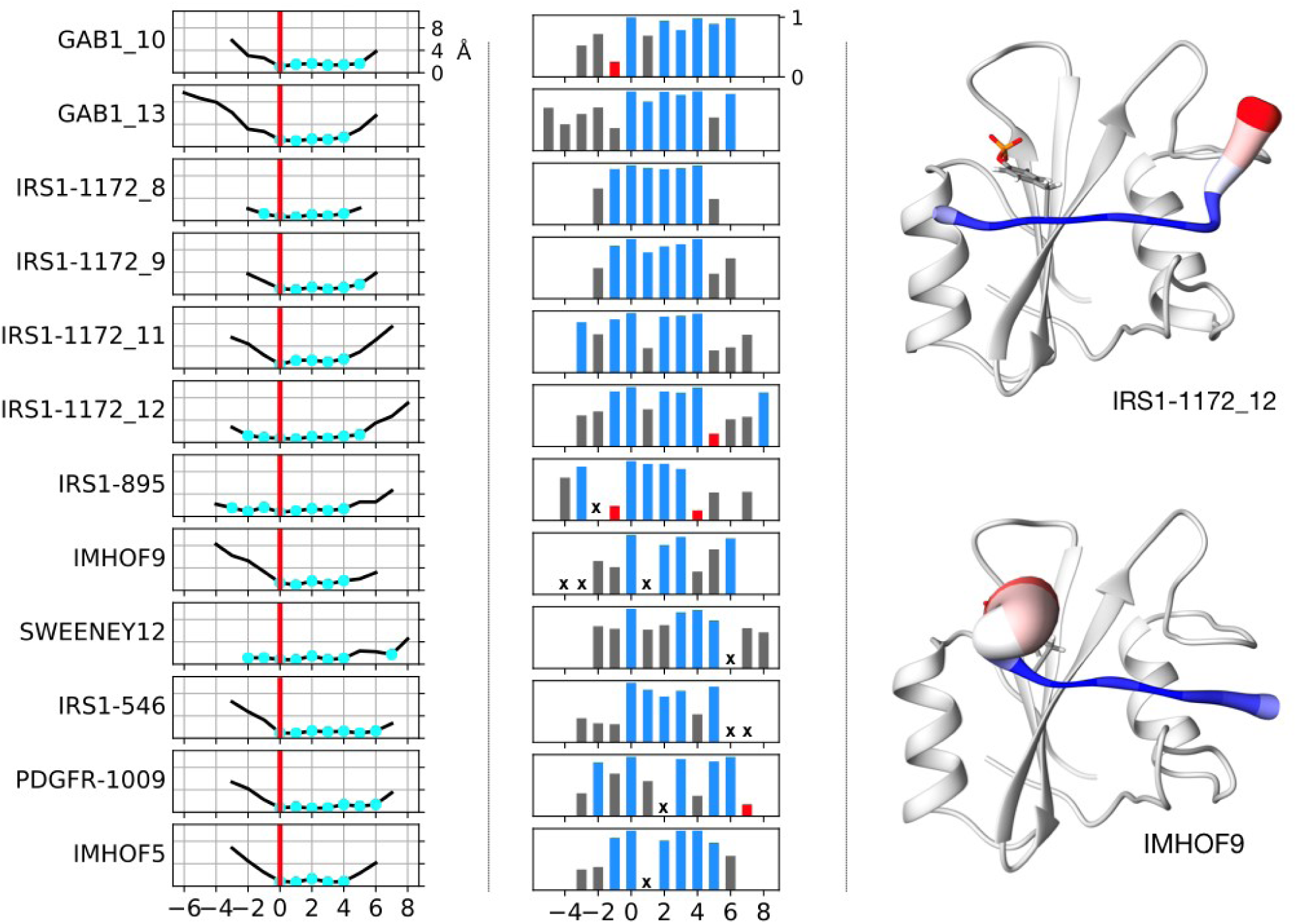
Dynamics of bound peptides. Left panel: RMSF of peptides bound to N-SH2. Residues whose RMSF is less then 1 Å larger than the minimal value are colored in cyan. Middle panel: side-chain order parameter Θ. Values close to unity indicate very narrow dihedral angle distributions and therefore bonds that are rigid with respect to rotation. Bars are colored according to the following scheme: Θ lower than 0.25 (red), between 0.25 and 0.75 (grey), greater than 0.75 (blue). A bold “x” indicates residues for which the side-chain order parameter cannot be defined (glycines and alanines). Right panel: most representative structures of the IRS1-1172_12 and IMHOF9 simulations, with the peptide backbone size and color (from blue to red) assigned based on the mobility of each residue.

### The central region of the peptide is in an extended conformation

Figure 3 shows the Ramachandran plots of the peptide backbone in the X-ray structures and in the MD simulations, for residues −2 to +5. In all cases, the dihedral angles of the conformations populated by residues from 0 to +3 fall in the top-left region of the plot, indicating an extremely stable extended structure [Cantor, 1980]. Residues −1 and +4 are extended, too, in all crystallographic structures, but they are more mobile in the simulations, populating regions of the Ramachandran plot corresponding to helical conformations in some cases. Beyond the −1 to +4 region, the backbone conformation is variable.

**Figure 3.**
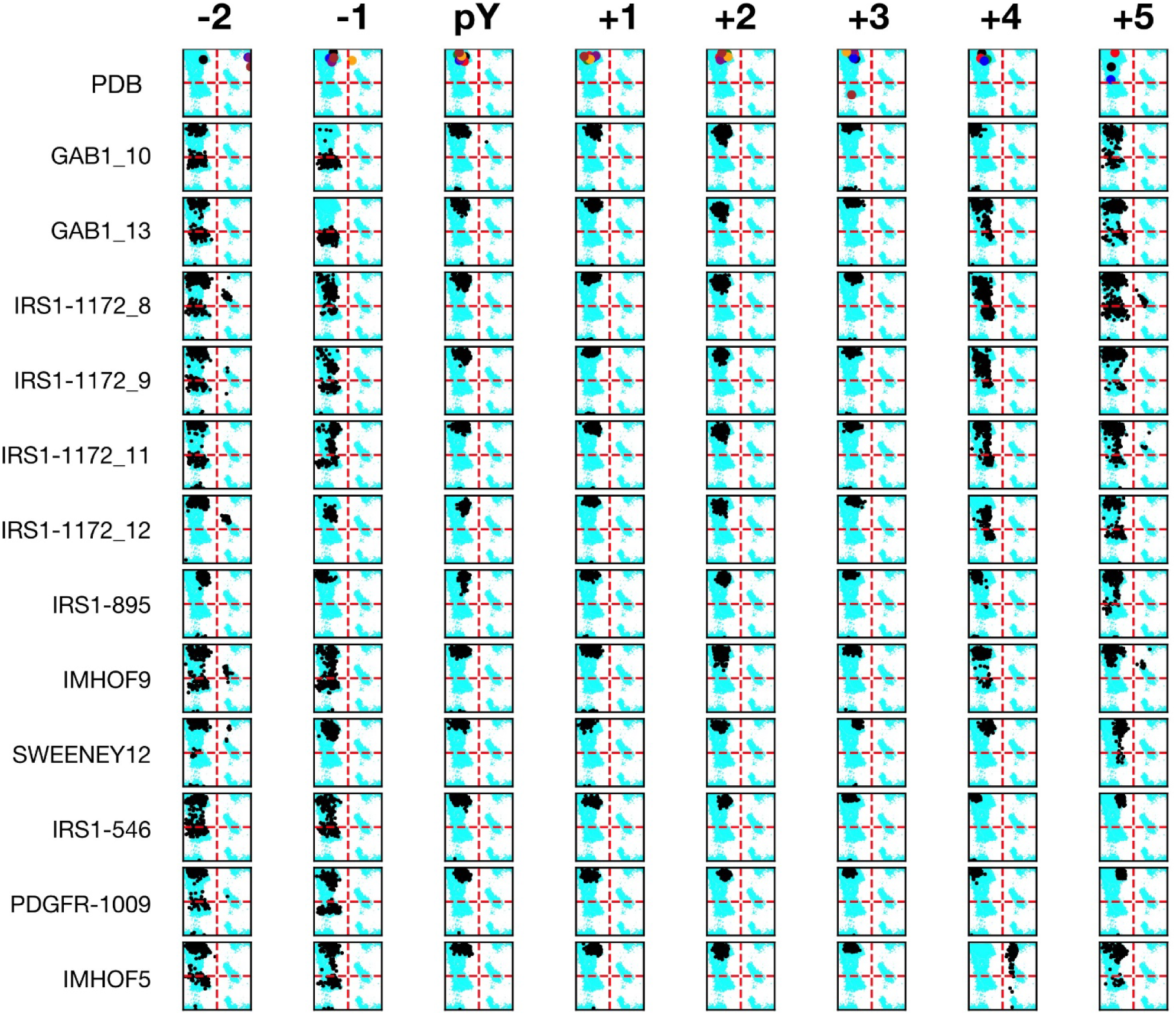
Backbone conformation of the bound peptides residues in PDB X-ray structures and in the simulations. Ramachandran plots of residues from position −2 to +5 with respect to pY are shown. Crystallographic structures are reported in the first line (“PDB”), with the following color code: 1AYA: green,1AYB: red, 3TL0: purple, 4QSY:black, 5DF6: orange, 5X7B: brown, 5X94: blue. The allowed regions of the Ramachandran plot are reported in cyan in the background. Angles ϕ and ψ are reported on the x and y axes, respectively, with values from −180° to +180°.

The extended peptide backbone conformation is stabilized by several H-bonds between the peptide and protein backbones, involving peptide residues −1, +1, +2 and +4, and protein residues H53 (βD4), K91 (BG7) and K89 (BG5), as illustrated in Figure 4. These interactions are present in some of the X-ray structures, and they are stably conserved in most of the MD simulations (Table 4). In addition, the MD trajectories show some transient interactions also for the backbone of residue +3 with K91 (BG7) and of +6 with Q86 (BG2) or G87 (BG3), which were not observed in the crystallographic structures.

**Figure 4.**
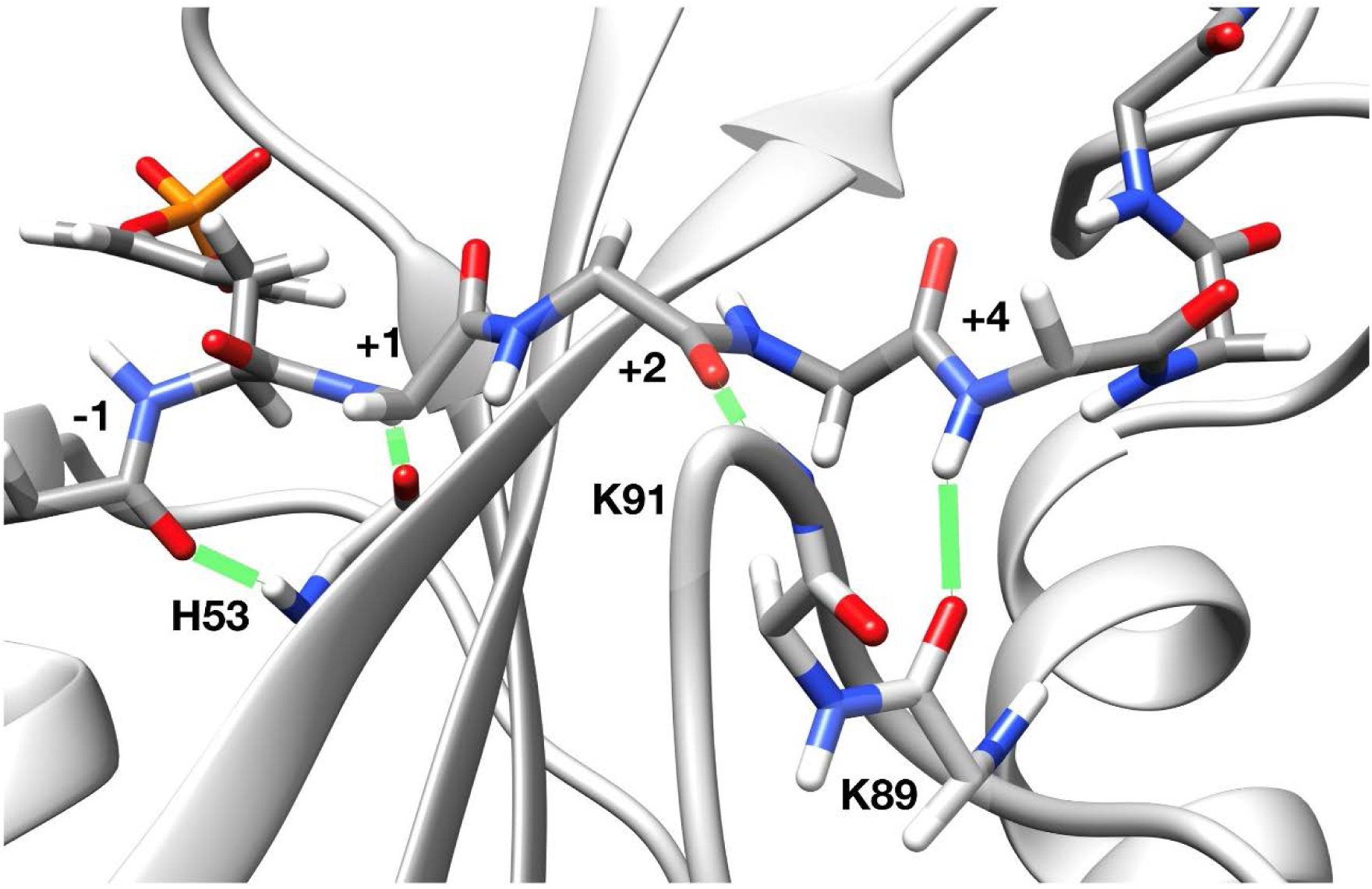
Main H-bonds between the peptide and protein backbones. Most representative structure of the IRS1-1172_12 simulation. H-bonds are highlighted by green lines.

**Table 4.**
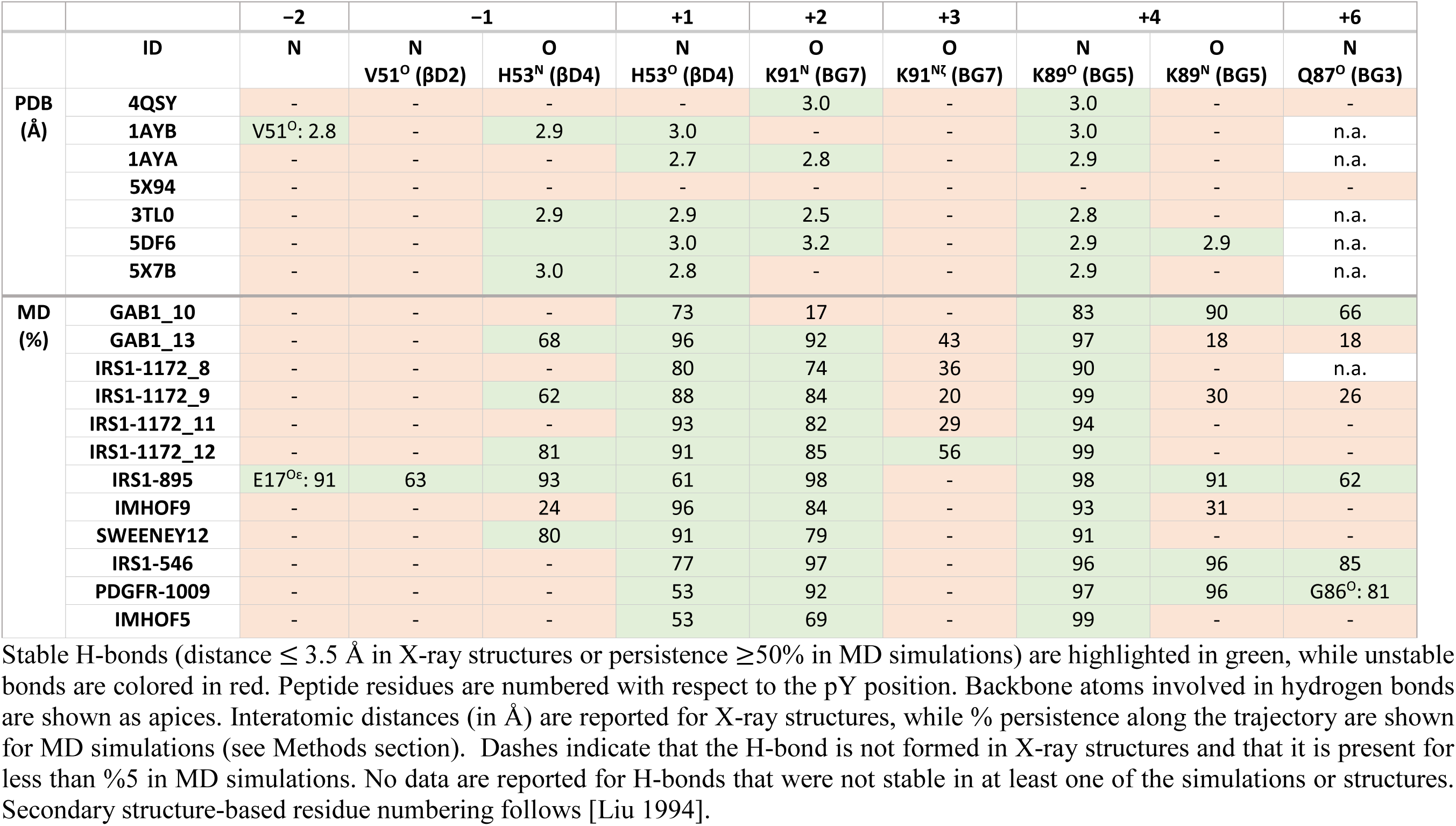
Hydrogen Bonds between the Peptide Backbone and the N-SH2 Domain.

### Phosphotyrosine interactions

The peptide position in the N-SH2 domain is strongly stabilized also by the interactions of the pY residue with its binding pocket. Several pY interactions are widely conserved in SH2 domains.

The most conserved residue is R βB5 (present in 98% of SH2 domains) [Liu 2012], which forms a salt bridge with the phosphate [Cohen 1995]. This is by far the most stabilizing interaction [Waksman 2004] and is responsible for the specificity for binding pY (as opposed to other phosphoamino acids): only the lengthy tyrosine side chain allows the phosphate to interact productively with this arginine, whereas serine and threonine are too short [Kuryian 1997, Mayer 1998].

R αA2 (present in 82% of SH2 domains [Liu 2012]) interacts with the phosphate group and makes an amino-aromatic interaction with the phenol ring of the pY [Bradshaw 2002].

K βD6 is located on the other side of the pY phenol ring from R αA2, so that the two residues together form a clamp around the pY [Bradshaw 2002].

The pY recognition site also contains an extensive network of hydrogen bonds [Cohen 1995]. In particular, S βB7 (present in 88% of SH2 domains) and T/S BC2 form direct hydrogen bonds with the phosphate. The BC loop backbone also contributes to H-bonding [Bradshaw 2002].

With respect to these general features of SH2 domains, the N-SH2 domain of SHP2 presents several peculiarities: it has a G in place of R αA2 [Waksman 2004] and in the crystallographic structures K βD6 contacts the phenol ring solely with its hydrocarbon chain and not with the amine [Cohen 1995].

Table 5 reports the H-bonds and the salt bridges formed by the phosphate in the crystallographic structures and in the simulations. The general picture described above is confirmed by our analysis of X-ray data. The R βB5 (R32)-pY phosphate ion pair is formed essentially in all structures, while K βD6 (K55) is always at a larger distance. H bonds with S βB7 (S34), S BC2 (S36) and the K BC1 (K35) backbone are always formed. An additional H-bond, present in all N-SH2 structures but not conserved in other SH2 domains, is formed with the side chain of T βC3 (T42).

**Table 5.** Hydrogen Bonds and Salt Bridges between pY and N-SH2 Residues.

|  | ID | Hydrogen bonds |  |  |  |  | Salt Bridges |  |  |
| --- | --- | --- | --- | --- | --- | --- | --- | --- | --- |
| | | S34<br>(βB7) | K35<br>(BC1) | S36<br>(BC2) | | T42<br>(βC3) | R32<br>(βB5) | K35<br>(BC1) | K55<br>(βD6) |
| | | Side-chain<br>Oγ | Backbone<br>N | Backbone<br>N | Side-chain<br>Oγ | Side-chain<br>Oγ1 | | | |
| **PDB**<br>(Å) | 4QSY | 2.7 | 2.6 | 3.0 | 2.7 | 2.8 | 4.5 | n.a. | 5.5 |
|  | 1AYB | 2.8 | 3.0 | 2.9 | 2.7 | 2.9 | 4.0 | n.a. | 4.5 |
|  | 1AYA | 2.5 | 2.8 | 2.9 | 2.4 | 2.5 | 4.0 | 7.9 | 4.8 |
|  | 5X94 | 3.2 | 3.2 | - | 2.5 | 3.4 | 4.5 | 9.0 | 6.1 |
|  | 3TL0 | 2.8 | 2.8 | 3.5 | 2.9 | 2.9 | 4.2 | n.a. | 4.7 |
|  | 5DF6 | 3.0 | 3.2 | 2.7 | 2.9 | 2.7 | 4.3 | n.a. | 6.9 |
|  | 5X7B | - | 2.8 | 2.6 | 3.3 | 3.2 | 4.0 | 6.6 | n.a. |
| **MD**<br>(%) | GAB1_10 | 100 | 95 | - | - | 99 | 100 | 79 | - |
|  | GAB1_13 | 98 | 94 | - | 27 | 99 | 100 | 62 | 65 |
|  | IRS1-1172_8 | - | - | - | - | 91 | 99 | - | 77 |
|  | IRS1-1172_9 | - | - | - | - | 38 | 32 | - | 94 |
|  | IRS1-1172_11 | 100 | 91 | 85 | 87 | 98 | 80 | - | 50 |
|  | IRS1-1172_12 | - | - | - | - | 96 | 99 | - | 82 |
|  | IRS1-895 | - | - | - | - | 84 | 91 | - | 75 |
|  | IMHOF9 | - | - | - | - | 61 | 83 | - | 69 |
|  | SWEENEY12 | 20 | - | - | - | 84 | 100 | - | 79 |
|  | IRS1-546 | 98 | 94 | 51 | 55 | 92 | 99 | 36 | 11 |
|  | PDGFR-1009 | - | - | - | - | - | 83 | - | 90 |
|  | IMHOF5 | - | - | - | - | - | 66 | - | 91 |

Stable bonds (distance $\leq 3.5$ Å for H-bonds and $\leq 4.0$ Å for salt bridges in X-ray structures or persistence $\geq 50\%$ in MD simulations) are highlighted in green, while unstable bonds are colored in red. Interatomic distances (in Å) are reported for X-ray structures, while % persistence along the trajectory are shown for MD simulations. Values lower than 5% are omitted. N.a. indicates X-ray structures where lysines 35 or 55 were not resolved in the electron density. Dashes indicate that the bond is not formed in X-ray structures and that it is present for less than %5 in MD simulations. Secondary structure-based residue numbering follows [Liu 1994].

In the MD simulations, the R βB5 (R32)–pY ion pair is stably maintained, as well as the H-bond formed by T βC3 (T42) (peculiar of SHP2 N-SH2). The other H bonds are less stable, indicating a significant mobility of the SH2 BC loop.

The distances between the pY phosphate and the charged side chains of R32, K35 and K55 are reported in Figure 5. Interestingly, the possible ion pair with K βD6 (K55), which is conserved in other SH2 domains, but surprisingly not present crystallographic structures of the N-SH2 domain [Lee 1994], does form often during the simulations. In addition, while the N-SH2 domain lacks the conserved R αA2, it has a K residue in position BC1 (K35), adjacent to the phosphate-binding site. In the crystallographic structures, its side chains points towards the solvent, but, in some of the simulations, conformational fluctuations of the BC loop allow the formation of this additional ion pair.

**Figure 5.**
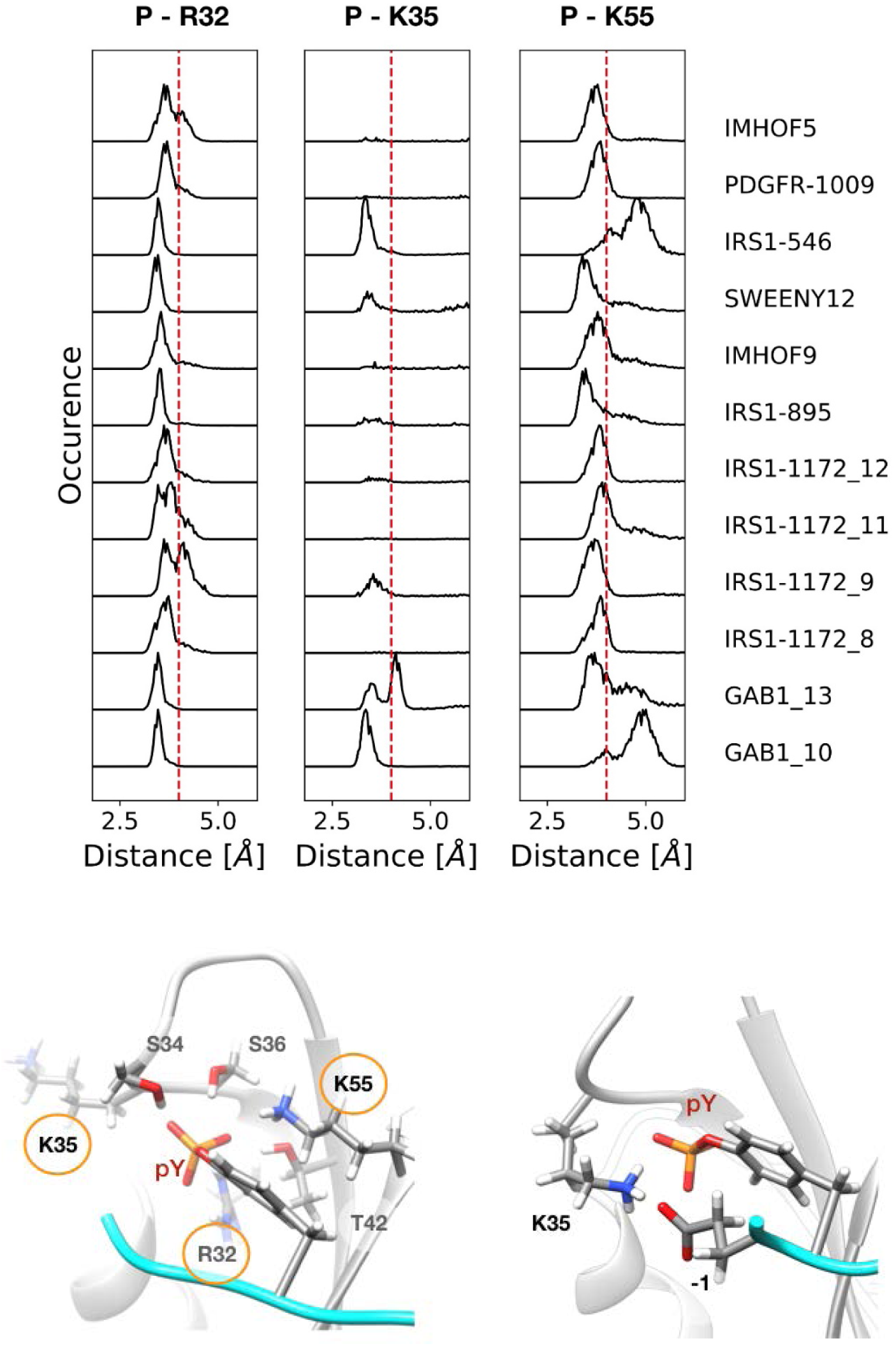
Most common ion-pair interactions between the pY phospahate and N-SH2 residues in MD trajectories. Top panel: distribution of distances between the phosphotyrosine phosphate and protein residues. Distances of less than 4 Å (vertical red dashed lines) are indicative of a stable salt-bridge. Bottom panels: N-SH2 residues that interacts with the phosphate group of pY (see Table 5) are shown on the left in the most representative structure of the IRS1-1172_8 simulation; the structure on the right shows the alternative arrangement of K35, where it interacts with the pY and a phosphopepeptide anionic residue in −1 (most representative structure of the GAB1_10 simulation).

Overall, the MD data suggest that notwithstanding a significant mobility of the pY pocket, a stabilization of the pY-domain interactions is mediated by an energetic compensation between H-bonds and salt bridges.

### Selectivity determining region: residues +1, +3 and +5 insert in hydrophobic pockets

Selectivity of SH2 domains is commonly considered to be determined mainly by residues C-terminal to the pY. Based on the interactions in this “selectivity determining region”, the domains have been classified in three classes [Yaffe 2002, Bradshaw 2002, Waksman 2004, Roque 2005]. The N-SH2 domain of SHP2 belongs to Type II, called open groove, or PLC-γ1-like, in which residues C-terminal to the pY bind in a long hydrophobic groove, delimited by EF and BG loops. This is illustrated in Figure 6, which shows the most representative conformation of the IRS1-1172_8 MD simulation. With the pY inserted in its binding pocket, the extended conformation of the peptide backbone forces residues +1, +3 and +5 to point towards the protein core and to insert into the hydrophobic ridge. Residues +2 and +4 point towards the solvent, but can interact with the loops BG and EF, which delimit the groove. In the N-terminal region of the peptide, residue −1 is solvent-exposed, while residue −2 points towards the protein surface, in the region of helix αA.

**Figure 6.**
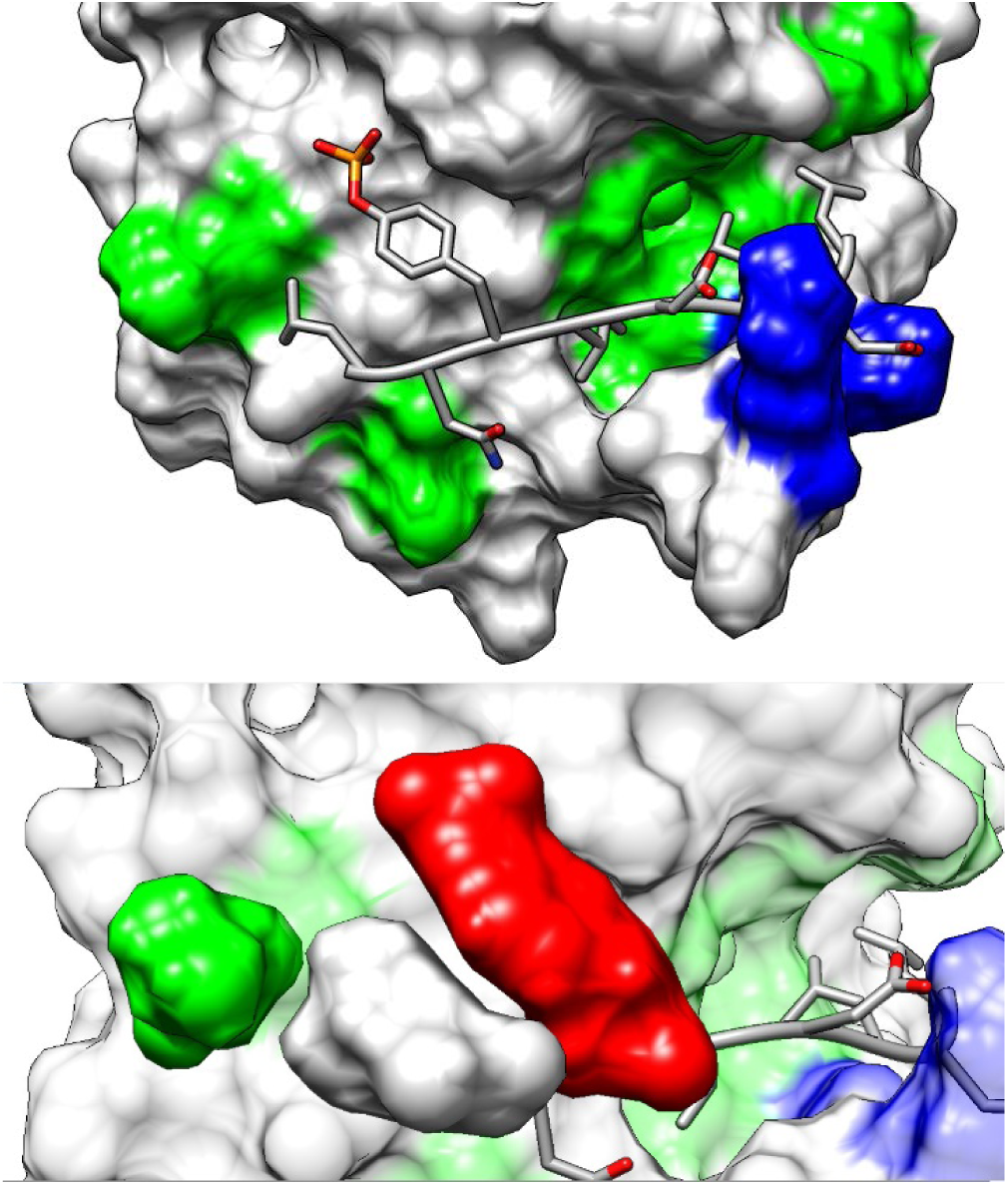
Most representative conformation in the IRS1-1172_8 MD simulation, illustrating the main specificity determining side chain interactions. Top: hydrophobic regions of the domain surface are shown in green, while cationic K89 and K91 are reported in blue. Bottom: interactions of the L−2 residue (grey surface), which inserts between the pY ring (red) and V14 (green).

Residues I+1, L+3 and L+5 form several interactions with hydrophobic amino acids that line the groove, remaining in contact with them for the whole length of the MD trajectory. In particular, residue +1 interacts with I54 (βD5), I96 (BG12) and methyl groups in the side chains of T52 (βD3) and E90 (BG6); residue +3 makes stable interactions with I54 (βD5), L65 (βE4), L88 (BG4) and I96 (BG12), and residue +5 with L65 (βE4), Y81 (αB9) and L88 (BG4). Interestingly, the +1 pocket is the only one where polar residues are present, in addition to hydrophobic amino acids. This might explain why peptide library studies and the sequences of known binding partners indicate that T can be present at position +1 of the peptide. As shown in Table 6, in the crystallographic structures 1AYA and 5DF6 (where a T is present in +1), no direct H-bond is formed between this residue and the protein domain. By contrast, our simulations show that an H bond can indeed be formed, either with T52 (βD3) or E90 (BG6). In one case (1AYA and PDGFR-1009) the peptide present in the crystal and in the simulation is the same one. However, the protein and peptide mobility, normally present in solution, can allow the formation of an H-bond that was not observed in the crystallographic structures.

**Table 6.** Hydrogen Bonds and Salt Bridges between Peptide Side Chains and the N-SH2 Domain.

|  | ID | H-bonds |  |  |  |  | Salt bridges |  |  |  |
| --- | --- | --- | --- | --- | --- | --- | --- | --- | --- | --- |
|  |  | -1 | +1 | +2 | +4 | +6 | -1<br>K35 (BC1) | +2<br>K91 (BG7) | +4<br>K89 (BG5) | +6<br>K91 (BG7) |
| P<br>D<br>B<br>Å | 4QSY | - | n.a. | - | - | D-Oδ<br>G68<sup>N</sup> (EF3): 3.0 | n.a. | D:7.6 | D:9.4 | D: 15 |
|  | 1AYB | - | n.a. | - | - | n.a. | n.a. | n.a. | E:7.3 | n.a. |
|  | 1AYA | n.a. | - | n.a. | - | n.a. | n.a. | n.a. | n.a. | n.a. |
|  | 5X94 | n.a. | n.a. | - | - | - | n.a. | n.a. | n.a. | n.a. |
|  | 3TL0 | - | n.a. | - | n.a. | n.a. | n.a. | n.a. | n.a. | n.a. |
|  | 5DF6 | - | - | - | - | n.a. | n.a. | E:4.2 | D:4.7 | n.a. |
|  | 5X7B | n.a. | n.a. | - | - | n.a. | n.a. | n.a. | n.a. | n.a. |
| M<br>D<br>% | GAB1_10 | * | n.a. | * | * | * | E:58 | D:62 | D:8 | D:23 |
|  | GAB1_13 | E-Oε<br>H53<sup>Nε</sup> (βD4): 42 | n.a. | * | * | * | E:23 | D:81 | D:16 | D:15 |
|  | IRS1-1172_8 | N-Nδ<br>V51<sup>O</sup> (βD2): 24 | n.a. | * | * | n.a. | n.a. | D:83 | D:16 | n.a. |
|  | IRS1-1172_9 | N-Nδ<br>V51<sup>O</sup> (βD2): 44 | n.a. | * | * | n.a. | n.a. | D:65 | D:22 | n.a. |
|  | IRS1-1172_11 | - | n.a. | * | * | n.a. | n.a. | D:71 | D:27 | n.a. |
|  | IRS1-1172_12 | N-Nδ<br>V51<sup>O</sup> (βD2): 19 | n.a. | * | * | n.a. | n.a. | D:83 | D:14 | n.a. |
|  | IRS1-895 | - | n.a. | - | E-Oε<br>N92<sup>N</sup> (BG8): 43 | n.a. | - | n.a. | E:30<br>E:18 (K91) | n.a. |
|  | IMHOF9 | - | n.a. | Q-Oε<br>K91<sup>Nζ</sup> (BG7): 21 | n.a. | n.a. | n.a. | n.a. | n.a. | n.a. |
|  | SWEENEY12 | n.a. | n.a. | Q-Oδ<br>K91<sup>Nζ</sup> (BG7): 29 | n.a. | n.a. | n.a. | n.a. | n.a. | n.a. |
|  | IRS1-546 | * | T-Oγ<br>T52<sup>Oγ</sup> (βD3): 24 | * | n.a. | n.a. | E:31 | E:57 | n.a. | n.a. |
|  | PDGFR-1009 | n.a. | T-Oε<br>E90<sup>Oγ</sup> (BG6): 49 | n.a. | Q-Nε<br>E90<sup>O</sup> (BG6): 22 | N-Nδ<br>Q87<sup>O</sup> (BG3): 62 | n.a. | n.a. | n.a. | n.a. |
|  | IMHOF5 | - | n.a. | Q-Oε<br>K91<sup>Nζ</sup> (BG7): 27 | W-Nε<br>Q87<sup>O</sup> (BG3): 91 | - | n.a. | n.a. | n.a. | n.a. |

To quantify the stability of the hydrophobic interactions between each peptide residue and the N-SH2 domain during all simulations, Table 7 reports the average values of the solvent accessible surface (SAS), for each side-chain. For comparison, the same parameter was calculated in the available crystallographic structures. For all the simulated sequences, residues +1 and +3 remain stably embedded in the domain grove. Residue +5 is also buried in all cases where a hydrophobic side chain is present at that position (with the single exception of the IRS1-1172_11 simulation).

**Table 7.**
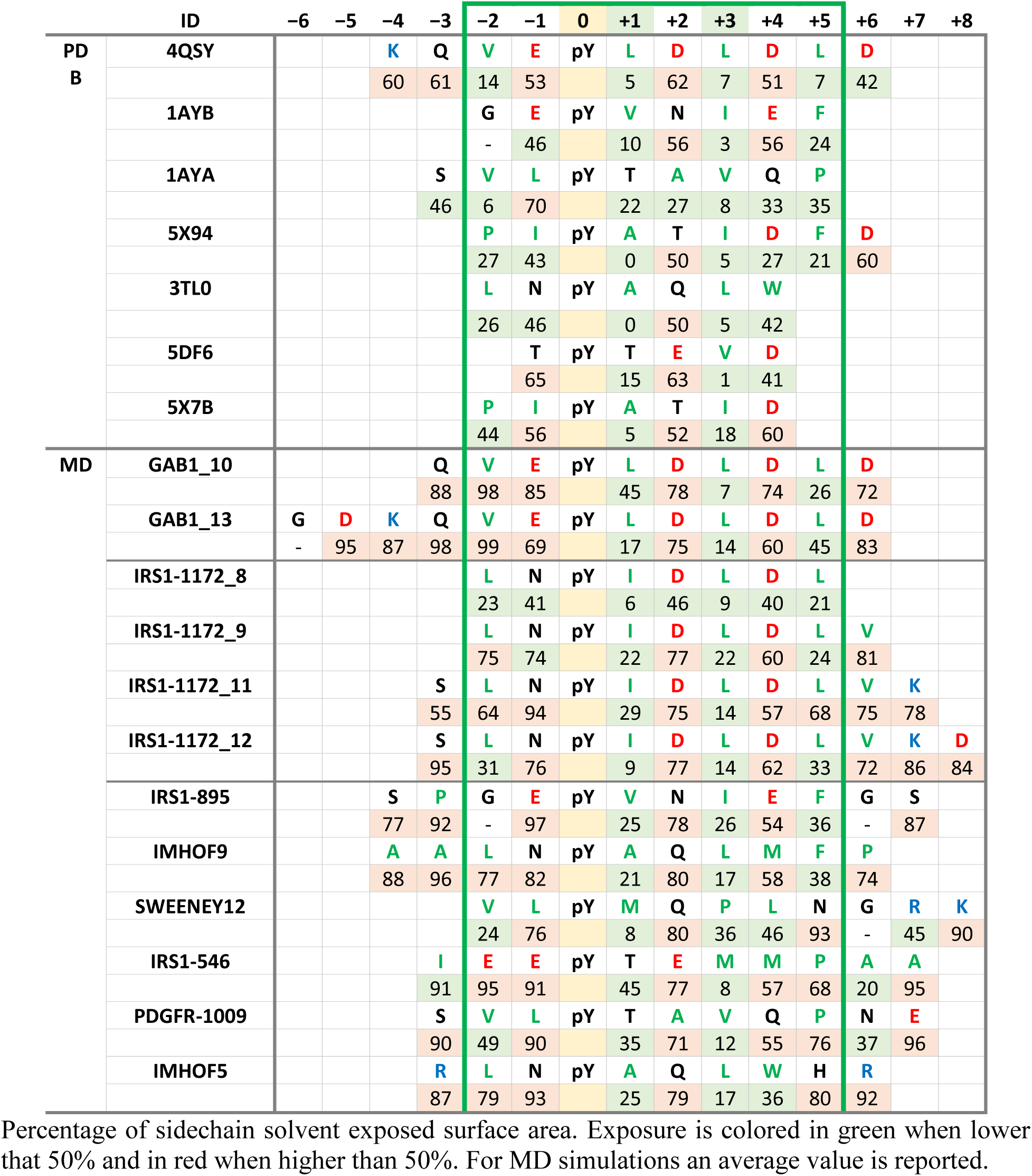
Solvent Exposure of Phosphopeptide Residues

Overall, the hydrophobic interactions involving residues +1, +3 and +5 of the peptide, which characterize type II SH2 domains, remained stable in most of our simulations, confirming their importance in determining the affinity and selectivity of the N-SH2 domain of SHP2.

#### Characteristic features of the SHP2 N-SH2 domain: interactions of residues −2, −1, +2 and +4

The N-SH2 domain of SHP2, while part of class II, presents peculiar features, which could affect its binding selectivity. As discussed in the section focusing on the pY interactions, more than 80% of SH2 domains have a conserved arginine at position αA2. By contrast, the SH2 domains of SHP2, SHP1 and MATK have a glycine at that position [Liu 2006]. In the N-SH2 domain, the lack of side chain at position 13 (G αA2) favors the accessibility of an exposed V14 at position αA3. This peculiarity has been previously described [Lee 1994; Bradshaw 2002; Waksman 2004], and explains why the N-SH2 of SHP2 is one of the few SH2 domains in which residues N-terminal to the pY contribute to the binding specificity. Indeed, in several simulations we observed that hydrophobic residues in −2 inserted between the pY ring and V14, interacting hydrophobically with both (Figure 6).

A second peculiarity, which has received limited attention in the literature, is that the N-SH2 domain has two K residues one amino acid apart in loop BG (K89 and K91, BG5 and BG7). The alignment of the human SH2 domains [Liu 2006] shows that positive residues in the BG loop are rather frequent. However, a (K/R-X-K/R) pattern is much less common, and is a rather unique characteristic of the SHP2 N-SH2 domain in the region of the loop facing toward the peptide binding groove. In principle, these side chains could form electrostatic interactions with acidic residues present in +2 and +4 of the peptide, which are shown to be favorable at those positions by peptide array studies and by the sequences of high affinity natural partners (Tables 1 and 2). In the available crystallographic structures, these interactions would be possible in 4QSY, 1AYB, 5X94, where a D/E residue is present at position +2, +4 or both. However, rather surprisingly, a bona fide ion pair is not formed in any of these structures (Table 6).

Among the simulated sequences, a D/E residue is present at position +2 or +4 (or both) in 8 of the 12 simulations. Differently from the X-ray conformations, our simulations show that a stable salt bridge forms between the +2 residue and K91 in all cases where this is possible. An ion pair between residues +4 and K89 forms, too, although only for a fraction of the simulation time (Table 6 and Figure 7). Interestingly, even polar, uncharged residues at positions +2 and +4 can interact with K89 and K91 by forming H bonds (which, again, were not observed in the crystallographic structures). Therefore, the simulations indicate that electrostatic or H-bonding interactions between the BG loop and residues +2 and +4 can contribute significantly to the binding affinity and selectivity.

**Figure 7.**
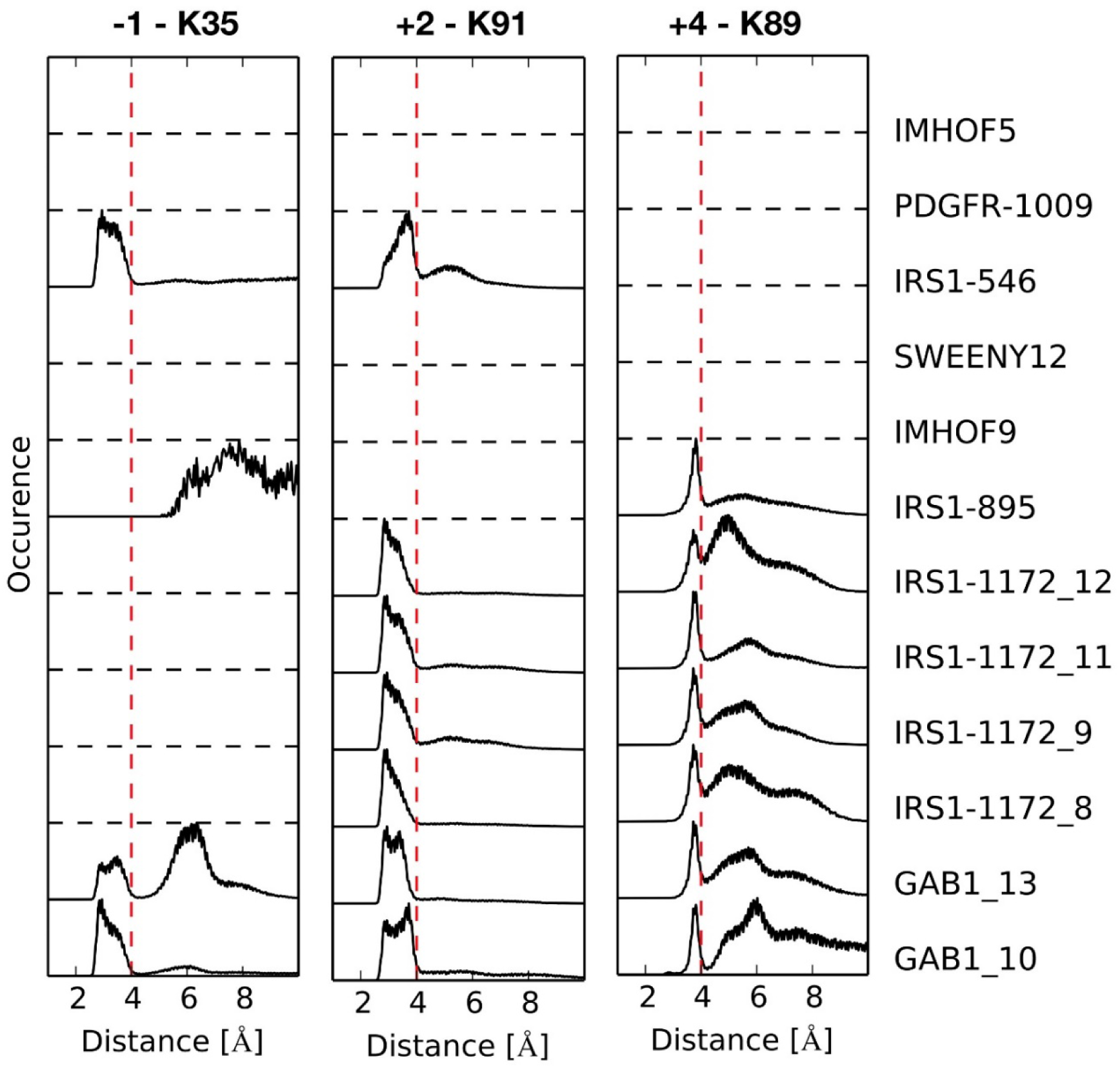
Most common intermolecular ion-pair interactions between the phosphopeptide side chains and the N-SH2 domain. Distribution of charged group distances populated in each MD trajectory. Distances of less than 4 Å (vertical red dashed lines) are indicative of a stable salt-bridge. Dashed horizontal lines indicate that the corresponding phosphopeptide sequences lack an anionic residue at those positions, and therefore cannot form the ion pair.

A third characteristic feature of the N-SH2 domain of SHP2 is the K residue at position BC1, which is present only in the C-terminal domain of ZAP70, while an R is present at that position in the N-SH2 domain of SHP1. As discussed above, in the crystallographic structures K35 always points towards the solvent. However, in the simulations, when E was present at position −1 (with the single exception of IRS-895), it interacted electrostatically with K35 (BC1). Interestingly, the trajectories in which this happened (GAB1 and IRS1-546) were the same in which the K35-pY ion pair was observed, as discussed above (Table 5). Probably, the negative residue in −1 favors a conformational transition which brings the K35 side chain from being solvent exposed to pointing towards the domain core, and in interaction with the pY, where it partially replaces K55 (Figure 5). A high mobility of K35 is supported by the observation that its side chain is not resolved in the electron density of several crystallographic structures (Table 5). In addition, during the simulations, polar residues in −1 could also form marginally stable H bonds, with amino acids of the βD strand.

Finally, our simulations showed that some interactions are possible also for negatively charged or polar residues in +6. An aspartate in that position can interact electrostatically with K91 (BG7) (although without forming a stable ion pair, due to the flexibility of the C-terminal end of the peptide). By contrast, in the crystallographic structure 4QSV D+6 and K 91 are very distant. In addition, the side chain of residue +6 can also form an H-bond with the EF or BG loops.

#### The N-SH2 domain populates different conformations

The data reported above on interactions in the pY binding pocket in the MD simulations indirectly suggested a significant conformational variability of this region. This is clearly shown by an overall analysis of the domain mobility in the 12 trajectories. As shown in Figure 8, the most mobile regions were the BC loop, which forms the pY pocket, and the EF and BG loops, which control access to the hydrophobic specificity region.

**Figure 8.**
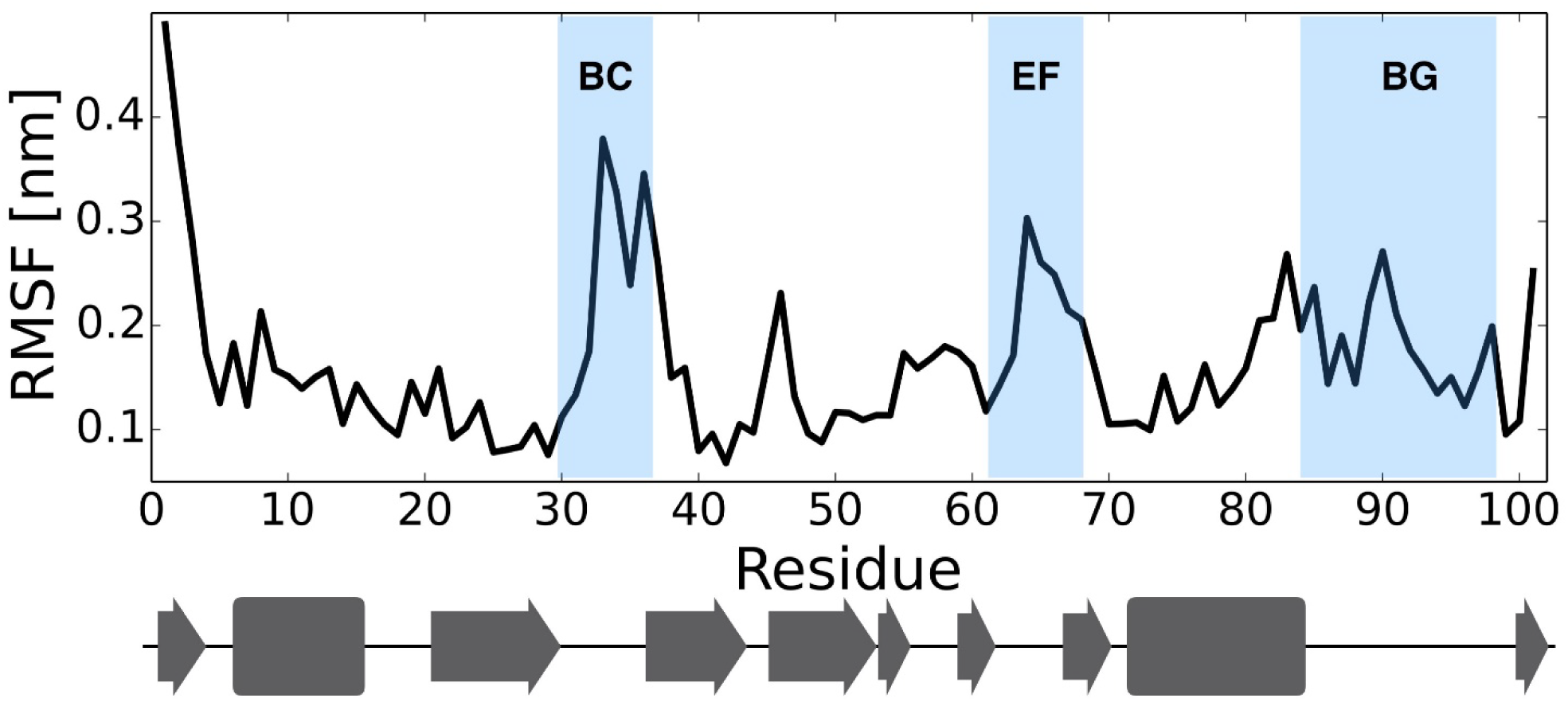
N-SH2 domain conformational variability in the 12 MD simulations. Root mean square fluctuations (RMSF) of the N-SH2 domain backbone in the cumulative trajectory including all 12 simulations. The domain secondary structure is reported at the bottom, for reference. The most mobile loops are highlighted in the figure.

Figure 9 analyzes the conformation of these flexible regions. For the BC loop it reports its average distance from T42, which is located in the pY pocket, on the βC strand (βC3) (Figure 9, left panel). While this loop is closed in all X-ray structures, in our simulations we find that it can change its structure significantly, populating also a more open conformation. Since this region is highly conserved in SH2 domains, we compared the MD conformations to those observed in experimental structures (both crystallographic and NMR, obtained in solution) of other SH2 domains. An open conformation of the BC loop is observed in only a few of the crystallographic structures, but is significantly populated in solution according to NMR data. Therefore, our simulations might have observed a previously undescribed conformation of the pY loop of the N-SH2 domain, possibly disfavored by the crystal environment and by intermolecular crystallographic contacts.

**Figure 9.**
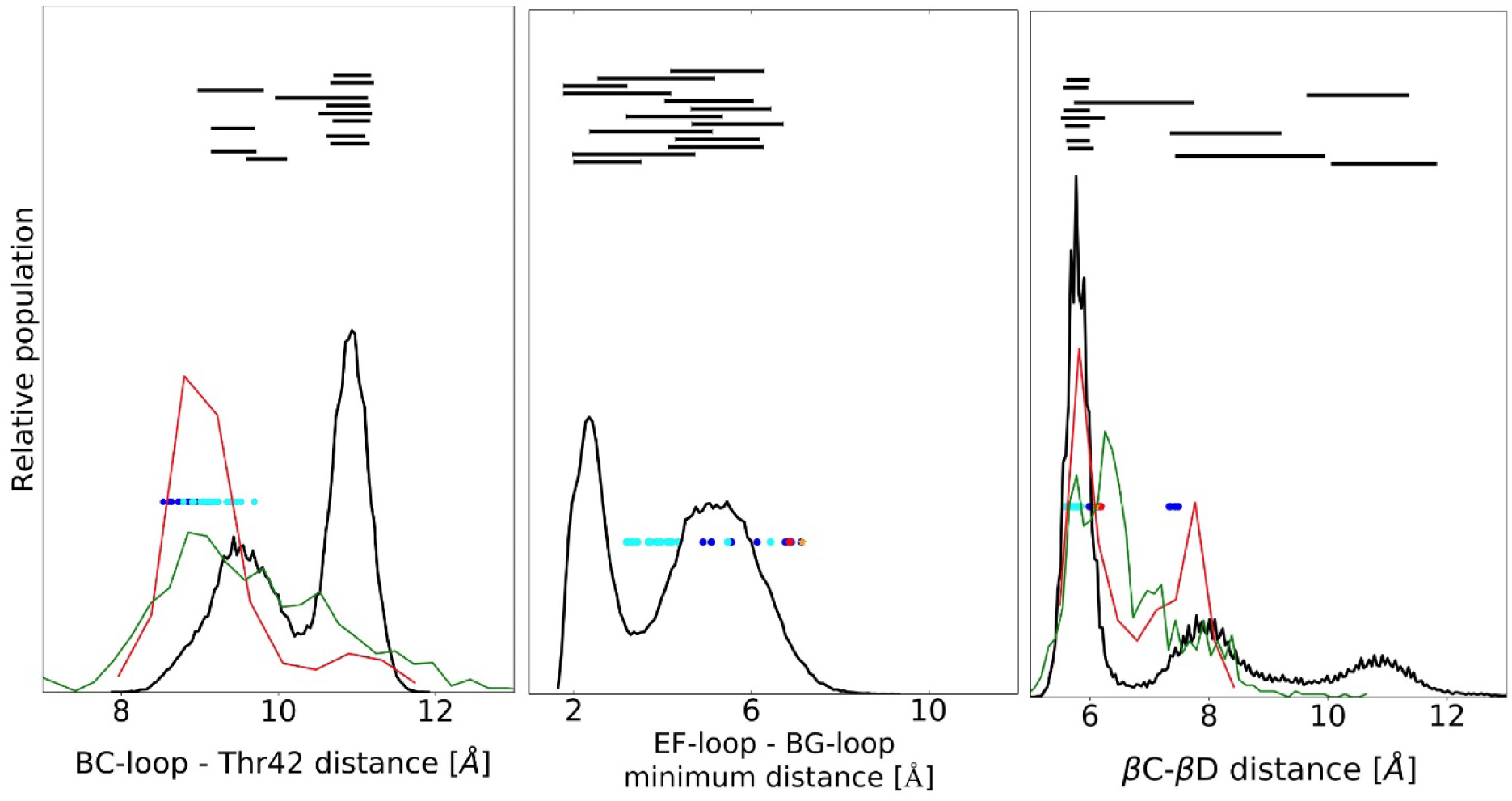
Structural parameters in simulated and experimental structures. Left: conformation of the pY pocket, as measured from the average distance between residues in the pY-loop (BC, residues 34−38) and T42 (βC3) in N-SH2, or structurally equivalent residues in other SH2 domains (see Sup. Mat.). Center: conformation of the loops controlling access to the selectivity determining region, as measured from the minimum distance between EF-loop (residues 66−69) and BG-loop (residues 84−96). Right: conformation of the central β sheet as measured from the inter-strand distance between C atom of D40 (βC1) and N atom of Q57 (βD’1) or structurally equivalent residues in other SH2 domains. Data from the overall MD simulation of 12 N-SH2:peptide complexes are shown in black, along with analogous data from X-ray (red) and NMR (green) structures of SH2 domains. Values for experimental structures of isolated N-SH2 domains are shown as blue (when phosphopeptide-bound) or red dots (with no bound peptide). Values for structures of the domain in the whole SHP2 protein are reported as cyan (autoinhibited conformation) or orange dots (active conformation). Average ± standard deviation of distances spanned by the individual simulations are indicated by black horizontal bars, reported in the order of Table 3, with Gab1_10 at the bottom and Imhof5 at the top.

The EF and BG loops, which regulate the accessibility of the specificity region, are distant in all structures of phosphopeptide/N-SH2 complexes, and more closed in the structure of the autoinhibited state of SHP2. Indeed, based on structural data, this transition has been hypothesized to be part of the allosteric switch controlling SHP2 activity and binding affinity [Barford 1998, Hof 1998, Darian 2011]. In our simulations, we find that the loops can close significantly even when a phosphopeptide is present in the binding cleft. In some cases, they close around the peptide, clasping it tightly and getting in contact. The high sequence variability of the BG loop does not allow a quantitative comparison with the structures of other SH2 domains, in this case. However, while such closed conformations have never been observed in X-ray structures of SHP2, for other SH2 domains the EF and BG loops have been described as a “set of jaws” that clamp down on the peptide [Eck 1993, Cohen 1995].

Another element of structural flexibility that we observed in our simulations is a variable length for the central β-sheet. As shown in Figure 9, values going from ∼5 to ∼12 Å are populated for the distance between the N-terminal residue of the C strand (D40, βC1) and the opposite residue in strand D (Q57, βD’1). A similar variability (although in a smaller range) is present in the X-ray structures of the N-SH2 domain and also in the experimental structures of other SH2 domains. However, to the best of our knowledge, this important feature of conformational flexibility has not been previously discussed. These different conformational features are illustrated in Figure 10, which reports the most representative structures of two simulations.

**Figure 10.**
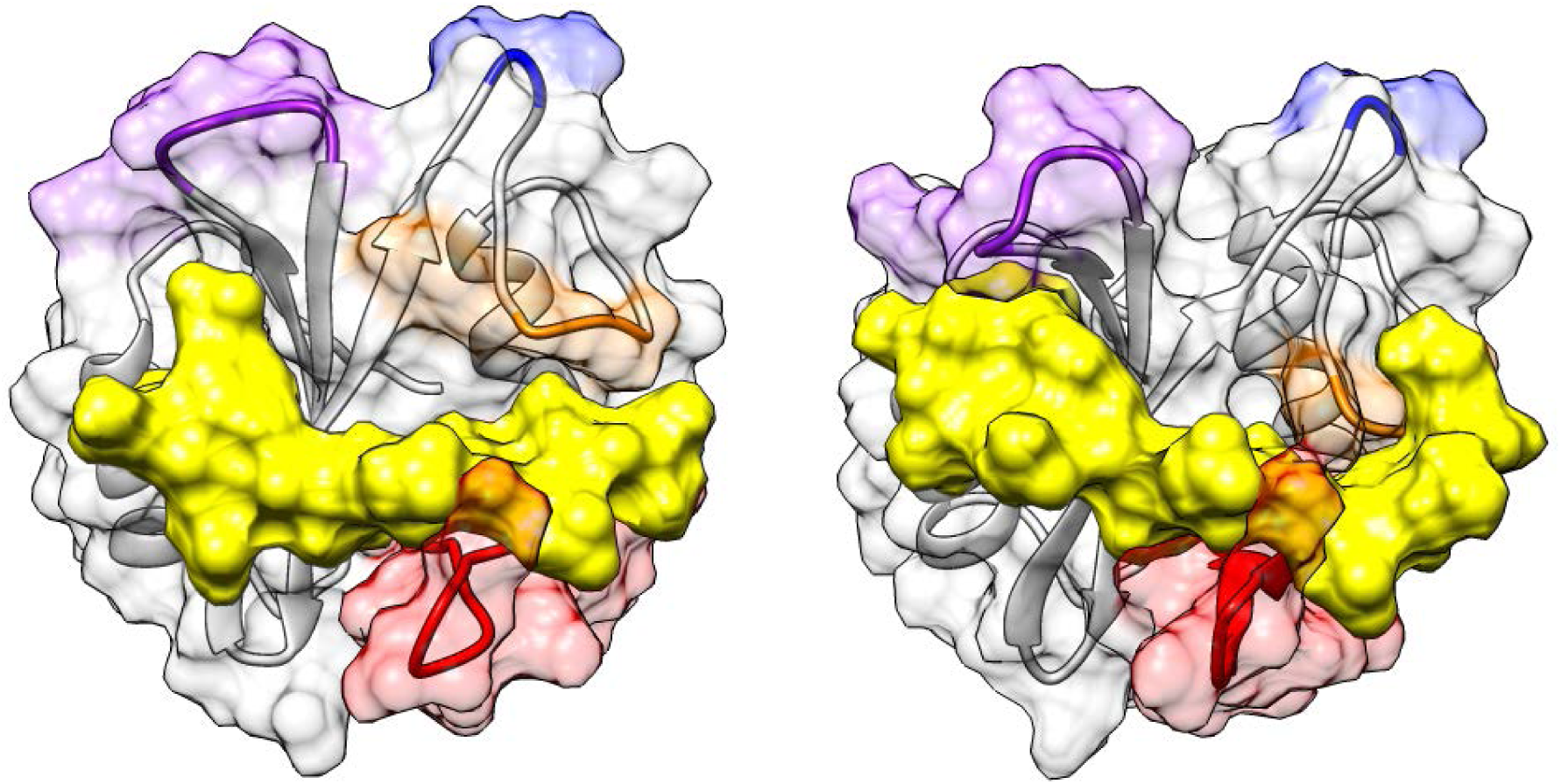
Conformational variability of the peptide-bound N-SH2 domain. Most representative structures of simulations IRS1-1172_9 and IRS1-1172_11, showing the conformational transitions of loops BC (purple), EF (orange) and BG (red) loops and of the central β sheet connecting strands C and D. The DE loop is highlighted in blue. The peptide surface is shown in yellow.

As shown in Figure 9, each individual simulation populated only one region of the overall conformational space. This finding could be due to an effect of the peptide sequence on the conformational properties of the domain, but it could also be caused by insufficient sampling of the conformational space in the single simulations. Further studies will be required to clarify these aspects.

## CONCLUSIONS

This work analyzed the structural determinants of the binding affinity and selectivity of the N-SH2 domain of SHP2. Some of the features responsible for the sequence preferences of this domain were already visible in the previously published crystallographic structures. The simulations confirmed that, even in solution and notwithstanding the significant motions of the domain and of the bound peptide, those interactions are conserved. In particular, residues −2 to +5 are stably interacting with the domain and this region of the peptide adopts an extended conformation (particularly from 0 to +3). The pY is stabilized in its pocket by multiple electrostatic and H-bonding interactions, while hydrophobic residues are needed at positions +1, +3 and +5, where they interact with apolar side chains of the domain binding groove.

These properties are common to Type II SH2 domains. However, the simulations confirmed some peculiarities of the N-SH2 domain of SHP2, which differentiate it from other SH2 domains and might contribute to its selectivity. Specifically, in place of the commonly conserved R αA2, the N-SH2 domain of SHP2 has G13. As a consequence, a hydrophobic peptide residue at position −2 can insert in the space left free by the missing side chain and interact with the accessible side chain of V14 αA3, as well as with the phenol ring of pY, stabilizing its orientation and the overall complex. Indeed, selectivity for residues N-terminal to the pY is peculiar of the N-SH2 domain. Another characteristic property of the N-SH2 domain of SHP2 is the non-conserved T42 in βC3, which forms a stable H-bond with the pY phosphate.

More importantly, the simulations highlighted some features that were not visible in the crystal structures, thus providing novel insights in the binding preferences of the N-SH2 domain. A peculiarity of this domain is the KXK motif in the region of the BG loop facing towards the peptide binding grove. Anionic residues at positions +2 and +4 strongly interact with the two K side chains. Even polar amino acids at those positions in the peptide sequence can interact with them through H-bonds. These observations are supported by the frequent presence of acidic residues at those positions in the sequences of natural binding partners, while a similar sequence selectivity had not emerged clearly from peptide library studies.

Another feature characterizing the N-SH2 domain is that, in some cases, interactions extended up to residue +6, through H-bond or ion pair formation with the EF or BG loops. This previously unexplored possibility warrants further investigations.

Polar amino acids at +1 can form H-bonds with residues in the corresponding domain pocket. This finding explains why a T residue was shown to be permitted at that position by library studies (in addition to hydrophobic amino acids).

Surprisingly, the conserved K βD6 does not form an ion pair with the pY phosphate in crystallographic structures. MD simulations indicated that domain motions actually allow significant electrostatic interactions with this residue in solution.

Another cationic residue is present in the pY pocket (K35, BC1), but in the crystallographic structures it points towards the solvent, without interacting with the pY. Our simulations showed that the presence of an acidic residue at position −1 of the phosphopeptide can favor a conformational transition that brings K35 towards the domain. In this new orientation, it interacts both with pY and with the residue in −1.

Finally, we observed in our simulations a significant conformational flexibility of the domain. These conformational transitions were associated with the BC loop (which forms the pY pocket), with the DE and BG loops controlling access to the peptide binding groove and with the central βC and βB strands, and were broader than those previously hypothesized, based on the different crystallographic structures of the domain. Investigation of the possible role of these motions in the function of SHP2 will require a more extensive exploration of the conformational properties of the N-SH2 domain.

## ASSOCIATED CONTENT

The following files are available free of charge.

Coordinates of the most representative structures of each MD trajectory (PDB format). Authors will release these coordinates upon article publication.

Supporting Information: methods used for the analysis of experimental SH2 domain structures (Figure 9) and Table S1 (preparation of the initial structures for the simulations.

## ACKNOWLEDGMENT

This work was supported by AIRC Foundation for Cancer Research in Italy (AIRC, grant IG19171, to L.S.), Italian Ministry of Education, University and Research (MIUR, grant PRIN 20157WW5EH_007, to L.S.), Fondazione Umberto Veronesi (Post-doctoral fellowship, to M.A.), Partnership for Advanced Computing in Europe (PRACE, grant 2019204928, to G. B.), which awarded computational resources at CINECA (Italy), and CINECA (grant HP10BL5G4C, to G. B.).

## ABBREVIATIONS

SHP2: SH2 domain-containing tyrosine phosphate
SH2: Src Homology 2 domain
pY: phosphotyrosine
MD: molecular dynamics
JMML: juvenile myelomonocytic leukemia
Erk: extracellular signal-regulated kinases
MAPK: Mitogen-activated protein kinase
RTK: Receptor tyrosine kinase
IRS-1: Insulin receptor substrate 1
PDGFR: Platelet-derived growth factor receptor
CagA: cytotoxin—associated gene A
Gab1: GRB2 associated binding protein
PLC-γ1: phospholipase C gamma 1
ZAP70: Zeta-chain-associated protein kinase 70
MOE: molecular operative environment
PME: particle-mesh Ewald
RMSF: root mean square fluctuations.

## TOC Graphics

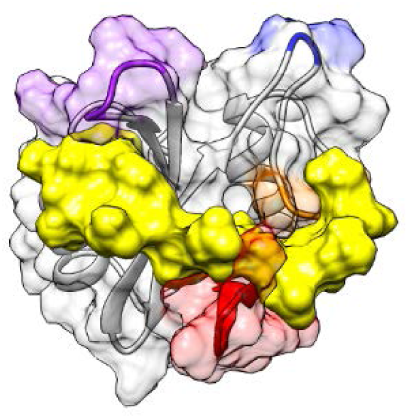

