## Supporting information for "Structural Determinants of Phosphopeptide Binding to the N-Terminal Src Homology 2 Domain of the SHP2 Phosphatase"

ANALYSIS OF AVAILABLE EXPERIMENTAL STRUCTURES OF SH2 DOMAINS

Two structural variables discussed for the MD simulations of the N-SH2 domain of SHP2 (Figure 9) were calculated also for the available experimental structures of SH2 domains. In the case of the N-SH2 domain, these variables were defined as:

- the "opening" of the central β-sheet, measured as the distance between $C_\alpha$ of residues Asp40 (strand β2) and Gln57 (strand β3);
- the "opening" of the pY-loop, measured as the average distance between $C_\alpha$ of Thr42 (strand β2) and $C_\alpha$ atoms of the five central residues belonging to the pY-loop (Ser34, Lys35, Ser36, Asn37, Pro38).

In the case of the experimental structures of SH2 domains, each structure was superimposed to 1AYD (X-ray structure of unbound SHP2 N-SH2), using the "matchmaker" function of UCSF Chimera [Pettersen 2004], and structurally equivalent residues were used to calculate the aforementioned variables.

The following experimental structures were used for this analysis:

**NMR structures:** 1ab2, 1aot, 1aou, 1bfi, 1bfj, 1blj, 1blk, 1csy, 1csz, 1fhs, 1fu5, 1fu6, 1ghu, 1hcs, 1hct, 1ju5, 1ka6, 1ka7, 1lui, 1luk, 1lum, 1lun, 1mw4, 1oo3, 1oo4, 1pic, 1qg1, 1rja, 1tce, 1wqu, 1x0n, 1x6c, 1z3k, 2bbu, 2cr4, 2crh, 2cs0, 2dcr, 2dly, 2dlz, 2dm0, 2dvj, 2ecd, 2ekx, 2el8, 2eo3, 2eo6, 2eob, 2etz, 2eu0, 2eyv, 2eyy, 2eyz, 2fci, 2ge9, 2gsb, 2jyq, 2k79, 2k7a, 2kk6, 2kno, 2l3t, 2l4k, 2l6k, 2lct, 2lnw, 2lnx, 2lqn, 2lqw, 2mc1, 2mk2, 2mqi, 2mrj, 2mrk, 2pld, 2ple, 2pna, 2pnb, 2rmx, 2ror, 2rsy, 2rvf, 2ysx, 2yu7, 3hck.

**X-ray structures:** 1a07, 1a08, 1a09, 1a1a, 1a1b, 1a1c, 1a1e, 1a81, 1ad5, 1aya, 1ayb, 1ayc, 1ayd, 1bf5, 1bg1, 1bhf, 1bhh, 1bkl, 1bkm, 1bm2, 1bmb, 1cj1, 1cwd, 1cwe, 1d1z, 1d4t, 1d4w, 1f1w, 1f2f,

1fbz, 1fmk, 1fyr, 1g83, 1gri, 1h9o, 1i3z, 1ijr, 1is0, 1jwo, 1jyq, 1jyr, 1jyu, 1k9a, 1kc2, 1ksw, 1lcj, 1lck, 1lkk, 1lkl, 1m27, 1m61, 1mil, 1nrv, 1nzl, 1nzv, 1o41, 1o42, 1o43, 1o44, 1o45, 1o46, 1o47, 1o48, 1o49, 1o4a, 1o4b, 1o4c, 1o4d, 1o4e, 1o4f, 1o4g, 1o4h, 1o4i, 1o4j, 1o4k, 1o4l, 1o4m, 1o4n, 1o4o, 1o4p, 1o4q, 1o4r, 1opk, 1opl, 1p13, 1qad, 1qcf, 1r1p, 1r1q, 1r1s, 1rpy, 1rqq, 1sha, 1shb, 1shd, 1skj, 1spr, 1sps, 1tze, 1uur, 1uus, 1x27, 1xa6, 1y1u, 1y57, 1yvl, 1zfp, 2abl, 2aoa, 2aob, 2aug, 2b3o, 2c0i, 2c0o, 2c0t, 2c9w, 2ci8, 2ci9, 2cia, 2dx0, 2fo0, 2h46, 2h5k, 2h8h, 2hck, 2hdv, 2hdx, 2hmh, 2huw, 2iug, 2iuh, 2iui, 2izv, 2oq1, 2ozo, 2ptk, 2qms, 2shp, 2src, 2vif, 2y3a, 3bkb, 3c7i, 3cbl, 3cd3, 3cwg, 3cxl, 3eac, 3eaz, 3gqi, 3gxw, 3gxx, 3hhm, 3hiz, 3imd, 3imj, 3in7, 3in8, 3k2m, 3kfj, 3m7f, 3maz, 3mxc, 3mxy, 3n7y, 3n84, 3n8m, 3nhn, 3ov1, 3ove, 3pjp, 3pqz, 3ps5, 3psj, 3psk, 3qwx, 3qwy, 3s8l, 3s8n, 3s8o, 3s9k, 3t04, 3tkz, 3tl0, 3uf4, 3us4, 3uyo, 3vrn, 3vro, 3vrp, 3vry, 3vrz, 3vs0, 3vs1, 3vs2, 3vs3, 3vs4, 3vs5, 3vs6, 3vs7, 3wa4, 4d8k, 4dgp, 4dgx, 4e68, 4e93, 4eih, 4ey0, 4f59, 4f5a, 4f5b, 4fbn, 4fl2, 4fl3, 4gl9, 4gwf, 4h1o, 4h34, 4je4, 4jeg, 4jgh, 4jmg, 4jmh, 4jps, 4k11, 4k2r, 4k44, 4k45, 4l1b, 4l23, 4l2y, 4lud, 4lue, 4m4z, 4nwf, 4nwg, 4ohd, 4ohe, 4ohh, 4ohi, 4ohl, 4ovu, 4ovv, 4p9v, 4p9z, 4qsy, 4roj, 4tzi, 4u17, 4u1p, 4u5w, 4waf, 4wwq, 4x6s, 4xey, 4xi2, 4xz0, 4xz1, 4y5u, 4y5w, 4ykn, 4z32, 4zop, 5aul, 5bo4, 5cdw, 5d0j, 5d39, 5dc0, 5dc4, 5dc9, 5df6, 5eel, 5eeq, 5eg3, 5ehp, 5ehr, 5fi4, 5i6v, 5ibm, 5ibs, 5itd, 6cmp, 6cmr, 6cms, 6crg

**Table S1. Simulations of complexes created starting from X-ray structures)**

The simulated sequence (2st column) was modeled from the original sequence (3rd column) as present in the respective PDB structure (PDB code in 4th column). Substitutions (red characters), were performed with Molecular Operative Environment (MOE), followed by conformational analysis, local energy minimization with side chain repacking. Italics indicates residues that were deleted from the original sequence.

| ID | Sequence | Original Sequence | PDB |
|---|---|---|---|
| **GAB1_10** | QVE-pY-LDLDLD | *GDK*QVEpYLDLDLD | 4qsy |
| **GAB1_13** | GDKQVE-pY-LDLDLD | GDKQVEpYLDLDLD | 4qsy |
| **IRS1-1172_8** | LN-pY-IDLDL | *GDKQ*VEpYLDLDL*D* | 4qsy |
| **IRS1-1171_9** | LN-pY-IDLDLV | *GDKQ*VEpYLDLDLD | 4qsy |
| **IRS1-1172_11** | SLN-pY-IDLDLVK | *GDK*QVEpYLDLDLD | 4qsy |
| **IRS1-1172_12** | SLN-pY-IDLDLVKD | *GDK*QVEpYLDLDLD | 4qsy |
| **IRS1-895** | PGE-pY-VNIEFGS | SPGEpYVNIEFGS | 1ayb |
| **IMHOF9** | AALN-pY-AQLMFP | SVLpYTAVQPNE | 1aya |
| **SWEENEY12** | VL-pY-MQPLNGRK | *S*VLpYTAVQPNE | 1aya |
| **IRS1-546** | IEE-pY-TEMMPAA | SVLpYTAVQPNE | 1aya |
| **PDGFR-1009** | SVL-pY-TAVQPNE | SVLpYTAVQPNE | 1aya |
| **IMHOF5** | RLN-pY-AQLWHR | RLNpYAQLWHR | 3tl0 |
